# Radiation-Induced Cellular Plasticity: A Strategy for Combatting Glioblastoma

**DOI:** 10.1101/2024.05.13.593985

**Authors:** Ling He, Daria Azizad, Kruttika Bhat, Angeliki Ioannidis, Carter J. Hoffmann, Evelyn Arambula, Aparna Bhaduri, Harley I. Kornblum, Frank Pajonk

**Affiliations:** Department of Radiation Oncology, David Geffen School of Medicine at UCLA; Jonsson Comprehensive Cancer Center at UCLA; Department of Biological Chemistry at UCLA; NPI-Semel Institute for Neuroscience & Human Behavior at UCLA; Department of Neurosurgery, David Geffen School of Medicine at UCLA

**Keywords:** Glioblastoma, radiation, forskolin, cAMP, scRNAseq, differentiation therapy, mouse model, survival

## Abstract

Glioblastoma is the deadliest brain cancer in adults and almost all patients succumb to the tumor. While surgery followed by chemo-radiotherapy significantly delays disease progression, these treatments do not lead to long-term tumor control and targeted therapies or biologics have so far failed to further improve survival.

Utilizing a transient radiation-induced state of multipotency we used the adenylcyclase activator forskolin to alter the cellular fate of glioma cells in response to radiation. The combined treatment induced the expression of neuronal markers in glioma cells, reduced proliferation and led to a distinct gene expression profile. scRNAseq revealed that the combined treatment forced glioma cells into a microglia- and neuron-like phenotypes. *In vivo* this treatment led to a loss of glioma stem cells and prolonged median survival in mouse models of glioblastoma. Collectively, our data suggest that revisiting a differentiation therapy with forskolin in combination with radiation could lead to clinical benefit.

## Introduction

Glioblastoma (GBM) is the deadliest brain tumor in adults. The current standard of care, surgery followed by chemo-radiotherapy has not changed in almost two decades and the median survival times of 15-18 months are unacceptably low. A large body of literature supports the hierarchical organization of GBM with glioma stem cells (GSCs) at the top of this hierarchy, able to regrow the tumor and to give rise to more differentiated GBM cells [1, 2]. Importantly, GSCs resist established anti-cancer therapies making them a main culprit in treatment failure [3, 4].

The debate over the identity of the cell of origin for GBM is not settled but neural stem cells (NSCs) and oligodendrocyte precursor cells (OPCs) are likely candidates [5] and GSCs share stem cell traits with these cell populations. While the stem cell/precursor state of NSCs and OPCs depends on the association with a supporting niche environment, GSCs utilize the same niche factors but also carry a mutational burden that supports stemness [6] and actively modulate pro-differentiation signaling to maintain GSCs traits and to block differentiation [7]. Overcoming such differentiation blocks in solid cancers has been attempted in the past. While numerous approaches have been successful *in vitro* [8–13], few succeeded *in vivo* [11, 12] and even fewer prolonged survival [12]. In GBM, this situation is further complicated by the requirement for systemically administered drug-based differentiation therapies to cross the blood-brain barrier (BBB).

We recently reported that ionizing radiation -aside from inducing cell death-led to global epigenetic changes in surviving cells with a transient acquisition of an open chromatin state in the promoter region of developmental transcription factors [14, 15]. Subsequently, some surviving non-stem cancer cells converted their phenotype into induced cancer stem cells [16, 17] and some surviving GSCs trans-differentiated into pericyte- and vascular-like cells [18], thus suggesting that exposure to ionizing radiation transiently elevates glioma cells into a multipotent state, comparable to lineage committed normal stem/progenitor cells.

Ideally, a differentiative therapy would direct cells towards a mitotically incompetent state. E.g., sarcoma cells can be forced into erythrocyte-like cells *in vitro* that even expel their nucleus [19]. In the CNS neurons are terminally differentiated cells that have lost their capacity to divide, thus making them a desirable end-state for a differentiation therapy in GBM. In the present study we sought to use this effect of ionizing radiation in combination with forskolin, an established agent for neuronal differentiation, to drive glioma cells into terminal differentiation.

## Materials and Methods

### Cell lines

Primary human glioma cell lines were established at UCLA as described in [1]; Characteristics of specific gliomasphere lines can be found in [20]. Primary GBM cells were propagated as gliomaspheres in serum-free conditions in ultra-low adhesion plates in DMEM/F12, supplemented with SM1 Neuronal Supplement (#05177, STEMCELL Technology, Kent, WA), EGF (#78006, STEMCELL Technology), bFGF (#78003, STEMCELL Technology) and heparin (1,000 USP Units/mL, NDC0409-2720-31, Lake Forest, IL) as described previously [1, 20, 21]. GL261 cells were cultured in log-growth phase in DMEM supplemented with 10% fetal bovine serum, penicillin, and streptomycin. All cells were grown in a humidified atmosphere at 37°C with 5% CO_2_. The unique identity of all patient-derived specimens was confirmed by DNA fingerprinting (Laragen, Culver City, CA). All lines were routinely tested for mycoplasma infection (#G238, Applied biological Materials, Ferndale, WA).

### Extreme Limiting Dilution Analysis (ELDA)

3x10^5^ HK-374 cells were intracranially implanted into the NSG mice as described above. Tumors were grown for 3 days for successful grafting. Tumor-bearing mice were then irradiated with a single dose of 4 Gy and injected intra-peritoneally on a 5-days on / 2-days off schedule for 2 weeks either with corn oil or forskolin starting 48 hours after the irradiation. The mice were then euthanized and tumor-bearing brains were dissected and further subjected for dissociation using mouse Tumor Dissociation Kit (Cat # 130-096-730, Miltenyi, Auburn, CA) to get single cell suspension, as described in [15]. The cells were counted and plated into the non-tissue-culture-treated 96-well plates at a range of 1 to 512 cells/well. Growth factors (EGF and bFGF) were supplemented every two days. Glioma spheres were counted 10 days later and presented as the percentage to the initial number of cells plated. The glioma stem cell frequency was calculated using the ELDA software [22].

### cAMP Assay

Primary GBM HK-374 monolayers were trypsinized and plated at a density of 5 x 10^4^ cells/well in a surfaced treated 96-well plate, while the HK374 gliomaspheres were dissociated and plated at the same density in a non-treated 96-well plate. The following day, cells were treated with freshly prepared Forskolin (#F3917, Sigma, St. Louis, MO) at 0.1, 0.25, 0.5, 1, 5, 10, 50, and 100 µM concentrations, with DMSO serving as the solvent control. 15 minutes after the treatment, the adherent monolayers were incubated with 100 µl/well cell lysis buffer from the cAMP Direct Immunoassays Kit (Fluorometric, ab138880, Abcam, Cambridge, UK), while the gliomaspheres were collected, centrifuged down and further incubated with 100 µl cell lysis buffer at RT for 10 minutes. 25 µl cell lysates were used to quantify the cAMP concentration by comparing to the standard cAMP curve, all the procedures were performed following the manufacturer’s guidelines.

### Quantitative Reverse Transcription-PCR

Total RNA was isolated using TRIZOL Reagent (Invitrogen, Waltham, MA). cDNA synthesis was carried out using the SuperScript Reverse Transcription IV (Invitrogen). Quantitative PCR was performed in the QuantStudio^TM^ 3 Real-Time PCR System (Applied Biosystems, Carlsbad, CA, USA) using the PowerUp^TM^ SYBR^TM^ Green Master Mix (Applied Biosystems). *C*_t_ for each gene was determined after normalization to PPIA and ΔΔ*C*_t_ was calculated relative to the designated reference sample. Gene expression values were then set equal to 2^−ΔΔCt^ as described by the manufacturer of the kit (Applied Biosystems). All PCR primers were synthesized by Invitrogen and used with PPIA as housekeeping gene (for primer sequences see **Suppl. Table 1**).

### Western Blotting

HK374 cells were plated and irradiated the next day with a single dose of 4 Gy. 48 hours after the irradiation, cells were daily treated with forskolin at 10 µM for 5 days. The cells were then lysed in 150 µl of ice-cold RIPA lysis buffer (10 mM Tris-HCl (pH 8.0), 1 mM EDTA, 1% Triton X-100, 0.1% Sodium Deoxycholate, 0.1% SDS, 140 mM NaCl, 1 mM PMSF) containing proteinase inhibitor (Thermo Fisher Scientific) and phosphatase inhibitor (Thermo Fisher Scientific). The protein concentration in each sample was determined by BCA protein assay (Thermo Fisher Scientific) and samples were denaturated in 4x Laemmli sample buffer (Bio-Rad) containing 10% β-mercaptoethanol for 10 minutes at 95 °C. Equal amounts of protein were loaded onto 10% SDS-PAGE gels and subjected to electrophoresis for 2 hours. Samples were then transferred onto 0.45 µm nitrocellulose membrane (Bio-Rad) and blocked in 1x TBST containing 5% bovine serum albumin (BSA) for 30 minutes at RT, followed by incubation with primary antibodies against Neurofilament-L (#2837S, 1:1000, Cell Signaling Technology), GFAP (#12389S, 1:1000, Cell Signaling Technology), β3-tubulin (#5568S, 1:1000, Cell Signaling Technology), and β-actin (#3700S, 1:1000, Cell Signaling Technology) in 1X TBST containing 5% BSA overnight at 4°C with gentle rocking. Membranes were then washed three times for 5 minutes each with 1X TBST and incubated with secondary antibodies, 1:5000 anti-mouse or anti-rabbit horseradish peroxidase (HRP; Cell Signaling) in TBST for two hours at RT with gentle rocking. Membranes were washed again three times for 5 minutes each with 1X TBST. Pierce ECL Plus Western Blotting Substrate (Thermo Fisher) was added to each membrane and incubated at RT for 5 minutes. The blots were then scanned with Odyssey Fc imaging system (LI-COR Biosciences, Lincoln, NE). β-actin was used as a loading control. Densitometry was performed using ImageJ. The ratio of the protein of interest over its endogenous control was calculated and expressed as relative intensity.

### Immunofluorescence

For *in vitro* neuron marker staining, HK374 cells were trypsinized and plated onto the round glass coverslips in 6-well plate at a density of 2 x 10^4^ cells/well. The following day, cells were irradiated at a single dose of 4 Gy and then daily treated with forskolin at 10 µM for 5 days starting from the 48 hours after irradiation. At day 5, the coverslips were fixed in formalin at RT for 15 minutes and washed three times with PBS, then permeabilized by 0.5% Triton X-100 for 10 minutes at RT. After three-time PBS washing, the coverslips were blocked with 10% goat serum diluted in PBS for 1 hour at RT and then incubated with primary antibodies against Neurofilament-L (Cell Signaling Technology, #2837, 1:100), and β3-tubulin (Cell Signaling Technology, #5568s, 1:400) overnight at 4°C. The next day, the secondary antibodies Alexa Fluor 594 Goat Anti-rabbit immunoglobulin G (IgG) (H/L) antibody (1:1,000 (Invitrogen)) or Alexa Fluor 488 Goat Anti-rabbit IgG (H/L) antibody (1:1,000 (Invitrogen)) were applied for 60 min, with subsequent nuclear counterstaining of Hoechst 33342 (Invitrogen, Cat# H3570, 1:5000). The sections were sealed with VECTASHIELD® PLUS Antifade Mounting Medium (Vector Laboratories, Cat#H-1900) and images were taken with a digital microscope (BZ-9000, Keyence, Itasca, IL).

For *in vivo* EGFP and neuron marker co-staining, brain sections were baked for 30 minutes in an oven at 65 °C, deparaffinized in two successive Xylene baths for 5 minutes each and then hydrated for 5 minutes each using an alcohol gradient (ethanol 100%, 90%, 70%, 50%, 25%). Antigen retrieval was performed using Heat Induced Epitope Retrieval in a citrate buffer (10 mM sodium citrate, 0.05% tween20, pH 6) with heating to 95 °C in a steamer for 20 minutes. After cooling down, the slides were blocked with 10% goat serum plus 1% BSA at room temperature for 30 minutes and then incubated with the primary antibodies against EGFP (Abcam, ab184601, 1:100) mixed with Neurofilament-L (Cell Signaling Technology, #2837, 1:100) or EGFP (Abcam, ab184601, 1:100) mixed with β3-tubulin (Cell Signaling Technology, #5568s, 1:400) overnight at 4°C. The secondary antibodies Alexa Fluor 594 Goat Anti-rabbit immunoglobulin G (IgG) (H/L) antibody (1:1,000 (Invitrogen)) and Alexa Fluor 488 Goat Anti-mouse IgG (H/L) antibody (1:1,000 (Invitrogen)) were applied followed by nuclear counterstaining and mounting procedures as above. Fluorescent images were then acquired using a confocal microscope (Nikon A1, Melville, NY).

### Animals

Female 6–8-week-old C57BL/6 mice, or NOD-*scid* IL2Rgamma^null^ (NSG) originally obtained from The Jackson Laboratories (Bar Harbor, ME) were re-derived, bred and maintained in a pathogen-free environment in the American Association of Laboratory Animal Care-accredited Animal Facilities of Department of Radiation Oncology, University of California, Los Angeles, in accordance with all local and national guidelines for the care of animals. Weight of the animals was recorded every day. 2x10^5^ GL261-Luc or 3x10^5^ HK-374-Luc cells were implanted into the right striatum of the brains of mice using a stereotactic frame (Kopf Instruments, Tujunga, CA) and a nano-injector pump (Stoelting, Wood Dale, IL). Injection coordinates were 0.5 mm anterior and 2.25 mm lateral to the bregma, at a depth of 3.0 mm from the surface of the brain. Tumors were grown for 3 days with successful grafting confirmed by bioluminescence imaging. Mice that lost 20% of their body weight or developed neurological deficits requiring euthanasia were sacrificed.

### Drug treatment

For *in vitro* studies, HK374, HK308 or HK157 cells were plated to form monolayers or gliomaspheres and irradiated at a single dose of 4 Gy the next day. 48 hours after irradiation, the cells were then treated with dibutyryl cAMP (dbcAMP; #D0627, Sigma), a cell-permeable analog of cyclic adenosine 3’5’-monophosphate (cAMP), at 1mM or forskolin at 10 µM for 5 consecutive days.

For *in vivo* studies, tumor grafting was confirmed via bioluminescence imaging. Mice implanted with either HK374 cells or GL261 cells were injected intraperitoneally with forskolin (#F3917, Sigma) at 5 mg/kg on a 5-days on / 2-days off schedule starting 48 hours after irradiation until they reached euthanasia endpoints. Forskolin was dissolved in corn oil containing 2.5% DMSO at a concentration of 0.55 mg/ml and prepared freshly for the injection.

### Irradiation

Cells were irradiated with at room temperature (RT) using an experimental X-ray irradiator (Gulmay Medical Inc. Atlanta, GA) at a dose rate of 5.519 Gy/min. Control samples were sham-irradiated. The X-ray beam was operated at 300 kV and hardened using a 4 mm Be, a 3 mm Al, and a 1.5 mm Cu filter and calibrated using NIST-traceable dosimetry. Corresponding controls were sham irradiated.

For *in vivo* irradiation experiments, mice were anesthetized prior to irradiation with an intra-peritoneal injection of 30 µL of a ketamine (100 mg/mL, Phoenix, MO) and xylazine (20 mg/mL, AnaSed, IL) mixture (4:1) and placed on their sides into an irradiation jig that allows for irradiation of the midbrain while shielding the esophagus, eyes, and the rest of the body. Animals received a single dose of 10 Gy on day 3 after tumor implantation.

### Cell cycle analysis

After forskolin (10 µM) treatment, HK374 cells were trypsinized and rinsed with ice-cold PBS. The cells were then centrifuged at 500 x g for 4 minutes, resuspended in 200 µl UltraPure RNase (Thermo Fisher, #12-091-021) and transferred to FACS tubes. 200 µl Propidium Iodide solution (1 mg/ml, Thermo Fisher, #P1304MP) was added and incubated for 15 minutes in the dark at RT. At least 100,000 events were analyzed by flow cytometry (LSR Fortessa, BD, San Jose, CA) and analyzed in FlowJo v10.

### Bulk RNA sequencing

For bulk RNA sequencing HK-374 cells were seeded into 6-well plates as monolayer cultures at 40k cells/well. 4 days after seeding the cells were irradiated with 4 Gy. Controls were sham irradiated. 48 hours later cells were treated with either forskolin (10 µM) or DMSO for 5 consecutive days. Total RNA was harvested and isolated using Trizol. Bulk RNA sequencing (RNAseq) was performed by Novogene and reads were mapped to the human genome (hg38) following their standard pipeline [15]. Read counts were analyzed using the iDEP package (version 2.0) [23]. Differentially expressed genes were calculated using the DESeq2 algorithm with a minimum of a 2-fold change and a false discovery rate (FDR) of 0.1. Enrichment *p*-values were calculated based on a one-sided hypergeometric test. *P*-values were then adjusted for multiple testing using the Benjamini-Hochberg procedure and converted to FDR. Fold Enrichment was defined as the percentage of genes in the list belonging to a pathway, divided by the corresponding percentage in the background.

### Single Cell RNA Sequencing

HK374 gliomaspheres were plated onto the Poly-D-Lysine/Laminin 6-well plate (#354595, Corning) and irradiated at a single dose of 4 Gy the next day. 48 hours after the irradiation, the cells were treated with forskolin at the concentration of 10 µM daily for 5 consecutive days or 21 days. At day 5 or day 21, the gliomaspheres in the suspension culture will be collected and dissociated with TrypLE (no phenol red, Thermo Fisher Scientific), and the adherent differentiated cells were washed with HBSS and de-attached with Trypsin/EDTA (Cell Applications, Cat#090K). The cells were then pooled and filtered through a 40-µm strainer and fixed with Evercode^TM^ Cell Fixation v2 kit (#ECF2101, Parse Biosciences, Seattle, WA) following the manufacturer’s guidelines. The samples (n=3 for each condition) were then sent out to the Genomics High Throughput Facility (GHTF) at the University of California, Irvine for subsequent single cell RNA sequencing (scRNAseq) using the Evercode^TM^ Whole Transcriptome Mini kit (#EC-W01010, Parse Biosciences). Sequencing reads from the mRNA libraries were mapped to the human genome (hg38) using the Parse Biosciences pipeline (split-pipe vers. 1.1.1) to generate cell by gene counts matrices. Data analysis was performed using the R package Seurat (version 4.3.3). Matrices were filtered for cells with high mitochondrial and ribosomal gene count and doublets were removed using DoubletFinder R package. For the subsequent cluster annotation, we used three published gene sets associated with cell types in the developing brain as previously described [24]. The trajectory analysis was executed using the Python package scVelo (version 0.2.5) [25]. Engagement of transcription factors was determined using the R package BITFAM [26].

### Statistics

Unless stated otherwise all data shown are represented as mean ± standard error mean (SEM) of at least 3 biologically independent experiments. A *p*-value of ≤0.05 in an unpaired two-sided *t*-test or one-sided ANOVA for multiple testing indicated a statistically significant difference. Kaplan-Meier estimates were calculated using the GraphPad Prism Software package (version 10.2.0). For Kaplan-Meier estimates a *p*-value of 0.05 in a log-rank test indicated a statistically significant difference. For bulk RNAseq, differentially expressed genes were calculated with a minimum fold-change of 2 and a false discovery rate cut-off of 0.1.

## Results

### Treatment of irradiated glioma cells with cAMP leads to expression of neuronal markers

In a first set of experiments, we tested the hypothesis that radiation induced a state multipotency that could be used to direct glioma cells into a neuron-like state using dbcAMP, part of established differentiation protocols for the neuronal differentiation of iPS cells [27, 28]. Patient-derived HK-374 glioma cells were irradiated with 0 or 4 Gy. We had previously shown that 48 hours after a single dose of 4 Gy, epigenetic remodeling led to a multipotent state with gains in open chromatin in the promoter regions of developmental transcription factors [15]. Therefore, we treated irradiated and unirradiated cells with dbcAMP (daily, 1 mM), starting 48 hours after irradiation **(Figure 1A)**. As early as 24 hours after start of the dbcAMP treatment both, dbcAMP-treated unirradiated and irradiated cells, showed elongated cell bodies reminiscent of neuron-like morphology (**Figure 1B, black arrows**) while control cells, not treated with dbcAMP, retained the morphology of untreated cells (**Figure 1B**).

**Figure 1.**
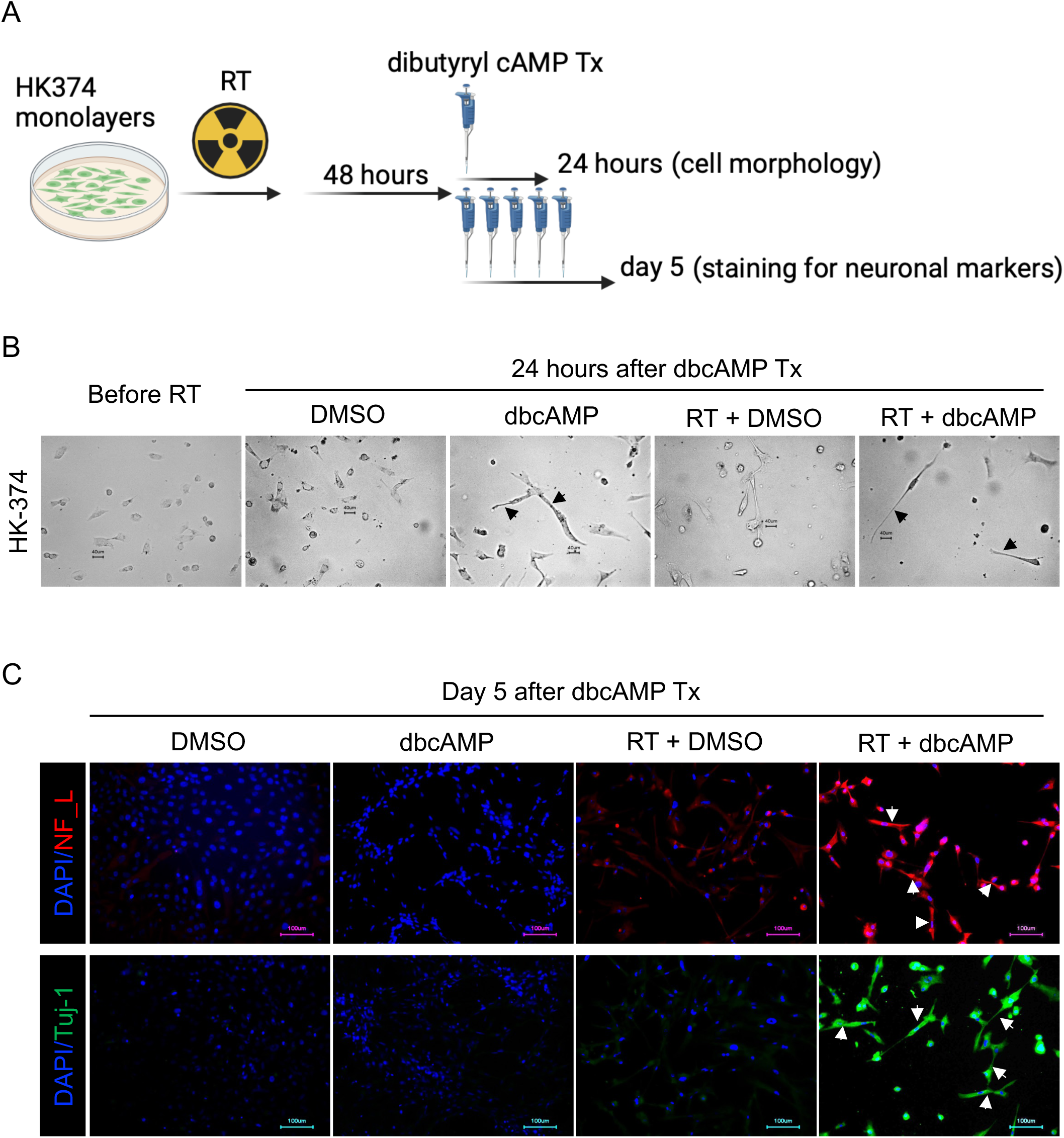
Neuronal marker induction by dbcAMP. Schematic of the experimental design for Figure 1 (**A**). Cells showed morphology changes into neuron-like cells (black arrows) as early as 24 hours after dbcAMP treatment start (**B**). The combination of radiation and dbcAMP but not radiation or dbcAMP treatment alone induced expression of neurofilament light chain (NF_L) and Tuj-1 (white arrows) (**C**). All experiments have been performed with at least 3 biological independent repeats.

After five daily treatments with dbcAMP, irradiated cells showed strong expression of the neuronal markers Tuj-1 and Neurofilament Light Chain **(Figure 1C, white arrows)**, which agreed with the neuron-like morphology of the cells. Cells treated with radiation or dbcAMP alone only showed faint expression of both markers (**Figure 1C**).

### The adenylate cyclase activator Forskolin induces neuron-like phenotype in irradiated glioma cells

*In vivo*, direct application of dbcAMP is not a feasible approach to differentiate glioma cells. To overcome this limitation, we next tested if forskolin, a known activator of adenylate cyclase [29] could increase cAMP levels in glioma cells. Glioma cells grown as monolayer cultures and gliomasphere cultures enriched for GSCs were treated with forskolin at concentrations of 0.1, 0.25, 0.5, 1, 5, 10, 50 or 100 µM. Differentiated cells and cells enriched for GSCs both showed a dose-dependent increase in cAMP production, thus confirming that forskolin can induce cAMP production in glioma cells, including GSCs. Induced intracellular cAMP concentrations did not reach mM concentrations used in dbcAMP experiments but plateaued below 200 nM after 15 minutes of forskolin treatment (**Figure 2A**).

**Figure 2.**
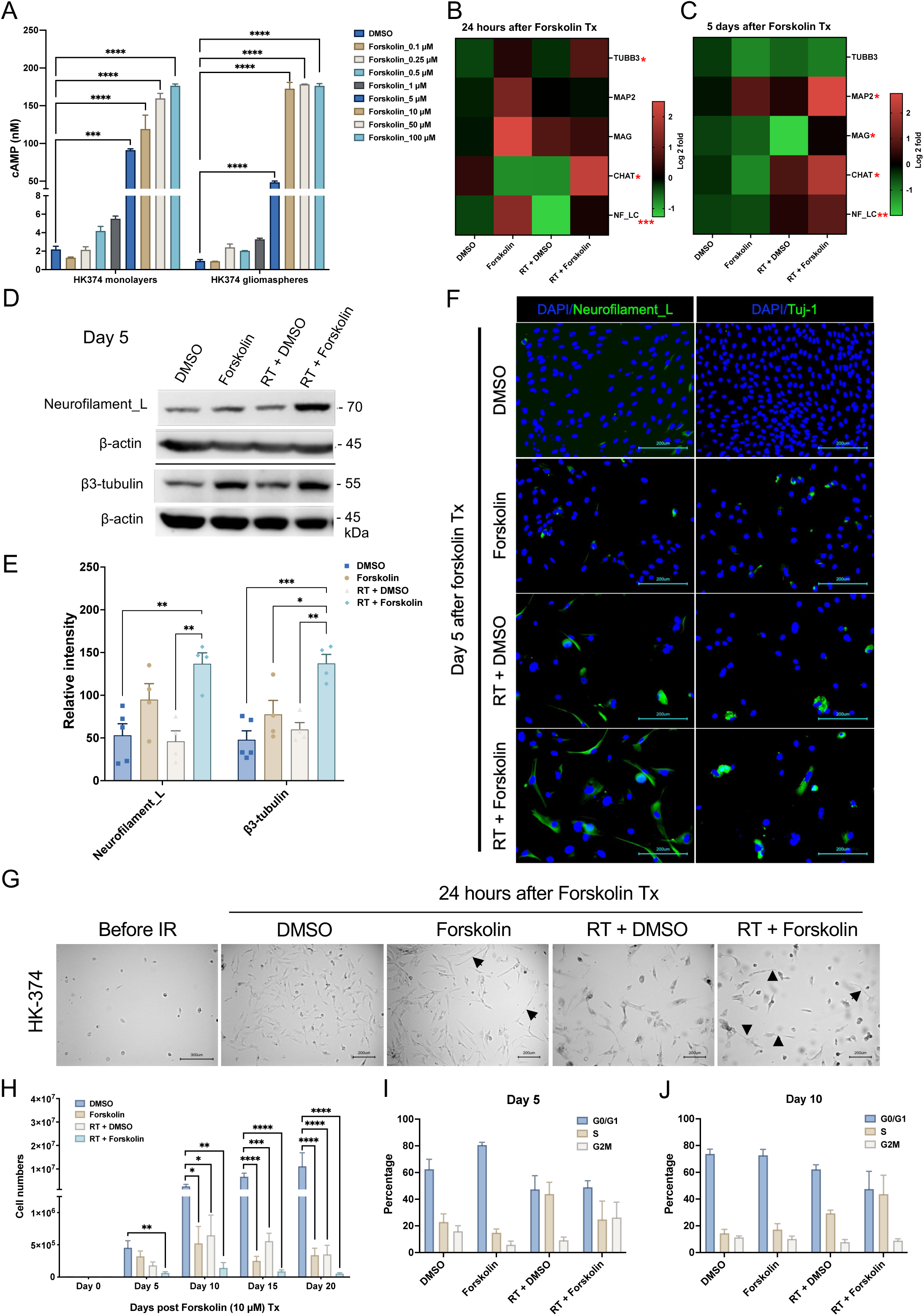
Forskolin effects on glioma cells. Treatment of HK-374 glioma monolayer cells and glioma spheres with forskolin leads to a dose-dependent induction of intracellular cAMP levels (**A**) and expression of neuronal markers at 24 h (**B**) and 5 days after start of forskolin treatment (**C)**. Representative Western blot for neurofilament light chain and β3-tubulin (**D**) shows a significant induction of both proteins after treatment with radiation and forskolin (**E**). Immunofluorescent imaging of glioma cells treated with radiation and forskolin shows expression of neurofilament light chain, and Tuj-1 in a subset of cells with elongated cell bodies (white arrows) (**F**). Morphology changes after irradiation and forskolin treatment could be observed as early as 24 hours after start of forskolin treatment (**G**, black arrows). Treatment of HK-374 glioma cells with radiation and consecutive treatment with forskolin for 5, 10 15 or 20 days significantly inhibits cell proliferation (**H**) and alters the cell cycle distribution (**I**, day 5; **J**, day 10). All experiments have been performed with at least 3 biological independent repeats. *p-*values were calculated using One-way ANOVA for **A, E, H**; The *p*-values listed in the heatmaps were from the comparison of RT + Forskolin to RT + DMSO. * *p*-value < 0.05, *\**\* *p*-value < 0.01, *** *p*-value < 0.001, **** *p*-value < 0.0001.

Next, we tested if treatment of irradiated glioma cells with forskolin would also induce the expression of neuronal markers. Quantitative RT-PCR 24 hours and 5 days after the start of forskolin treatment revealed that combined treatment with radiation and forskolin increased the expression of the neuronal markers β3-tubulin, CHAT, Neurofilament light chain (NF-LC), and MAG, thus indicating possible induction of a differentiated cell state (**Figure 2B/C**). We further confirmed the induction of neuronal markers NF-LC and β3-tubulin on day 5 after irradiation and forskolin treatment by western blotting (**Figure 2D/E)** and immunofluorescence staining **(Figure 2F, white arrows)**. However, expression of these markers was less uniform than after treatment with dbcAMP, consistent with much lower intracellular cAMP concentrations (**Figure 2A**). Forskolin alone or in combination with radiation induced changes in cell morphology with elongation of some of the cells’ bodies **(Figure 2G, black arrows)**. Considering the distinct characteristics of different TCGA subtypes in GBM, we included another two patient-derived GBM lines **(Suppl. Table 2)** to assess the effect of radiation plus forskolin on neuron-like differentiation. In HK-308 cells (mesenchymal subtype) combined treatment with radiation and forskolin showed similar induction of neuronal markers **(Suppl. Figure 1A)**, while these effects were less notable in HK-157 cells (proneural subtype) **(Suppl. Figure 1B)**.

### Forskolin in combination with radiation impacts cell proliferation and cell cycle in glioma cells

True differentiation towards a neuronal phenotype would result in a loss of mitotic capacity. To test the proliferative capacity of cells treated with radiation and/or forskolin, HK-374, HK-308 or HK-157 patient-derived glioma cells were seeded and treated with 0 or 4 Gy of radiation the following day. After an additional 48 hours, cells were treated with DMSO or forskolin (daily at 10 µM). Cell numbers were counted on day 5, 10, 15 and 20 after forskolin treatment initiation. While radiation or forskolin alone reduced cell numbers and/or slowed proliferation in all three lines to some extent, the combination of radiation and forskolin had additive effects on cell numbers from day 5 on (**Figure 2H, Suppl. Figure 2A/B**), thus suggesting that the combined effect is not cell line specific. Forskolin is known to affect cell cycle progression through specific inhibition of G1-to-S phase progression [30]. To study the lack of proliferation in more details we next performed a cell cycle analysis. On day 5, forskolin treatment led to an arrest in the G_1_/G_0_ phase of the cell cycle. Irradiation with a single dose of 4 Gy decreased the number of cells in the G_1_/G_0_- and G_2_/M-phase and increased the population of cells in S-phase, consistent with the well-known initial arrest at the G_1_/G_0_ and G_2_/M checkpoints and the subsequent release after DNA repair. Combined treatment with radiation and forskolin decreased the number of cells in G_1_/G_0_ while elevating the size of the S- and G_2_/M-phase cell populations (**Figure 2I)**.

On day 10, the G_1_/G_0_ population of untreated control cells increased, consistent with growth inhibition in confluent monolayer cultures. The cell cycle distribution of forskolin treated cells was similar to the distribution on day 5, while irradiated cells continued to redistribute, matching distributions of untreated control cells. Irradiated cells treated with forskolin appeared to be arrested at the G_1_-to-S transition of the cell cycle **(Figure 2J)**.

### Bulk RNA sequencing

After observing the effects of increased neuronal differentiation in GBM cells in response to radiation combined with forskolin treatment, in order to assess which pathways were activated in cells irradiated and subsequently treated with forskolin and if surviving cells would display gene expression profiles associated with neuronal differentiation, we performed bulk RNA sequencing on HK-374 cells cultured as monolayers. Compared to unirradiated controls, cells irradiated with 4 Gy had 421 differentially expressed genes (DEGs) at least 2-fold (FDR 0.1) up- and 865 genes down-regulated. After combined treatment we found 1,471 genes differentially up- and 2,058 genes down-regulated when compared to untreated control cells. Compared to irradiated cells, we found 945 DEGs up- and 704 genes down-regulated in cells treated with radiation and forskolin (**Figure 3A/B**), thus leading to a distinct gene expression profile for cells treated with radiation and forskolin (**Figure 3C**).

**Figure 3.**
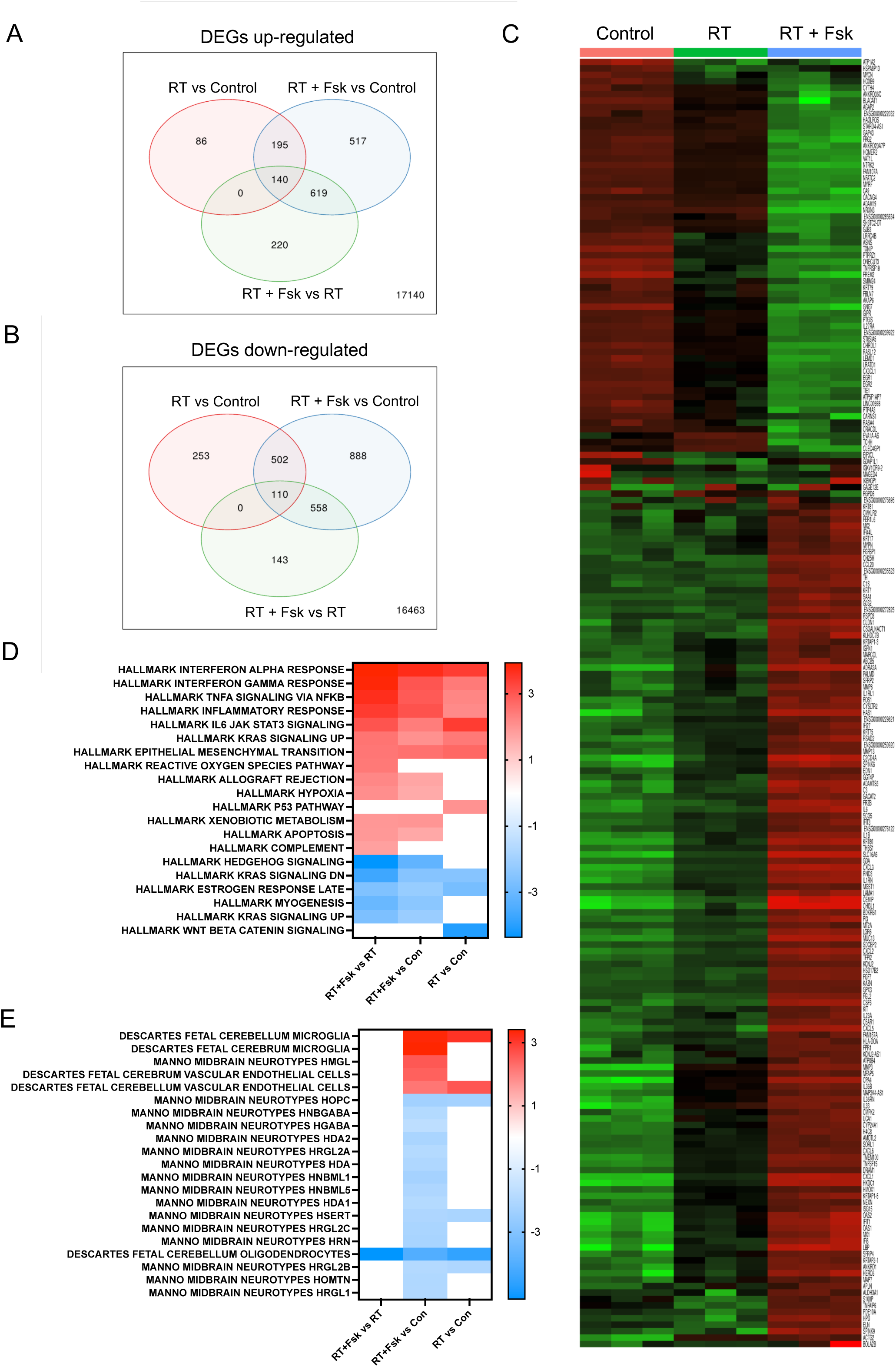
Bulk RNAseq of cells treated with radiation and forskolin. Treatment with radiation and forskolin induced the upregulation (**A**) of 220 and downregulation **(B)** of 143 unique differentially expressed genes leading to a distinctive gene expression profile. Heatmap showing the top 200 differentially expressed genes (**C**). Enrichment analysis using the Hallmark.MSigDB gene set and the differentially expressed genes indicated enhancement of proinflammatory signaling after treatment with radiation and forskolin when compared to radiation alone, induction of reactive oxygen species and inhibition of Shh and Kras signaling (**D**). Compared to unirradiated control cells the combination of radiation and forskolin led to gene expression profiles overlapping with those of microglia, neurons, and endothelial cells (Celltype.MSig.DB) (**E**). Abbrev. RT+Fsk represents RT + forskolin.

Enrichment analysis of upregulated DEGs found in irradiated samples compared to unirradiated control cells using the Hallmark.MSigDB gene set revealed overlap with genes associated with proinflammatory pathways, epithelial-to-mesenchymal transition, and activation of the p53-dependent response (**Figure 3D**), all consistent with the well-known effects of ionizing radiation [31]. The inflammatory response was amplified by the addition of forskolin and suppression of genes in Hedgehog signaling.

Enrichment analysis of upregulated DEGs in cells treated with radiation and forskolin and compared to control cells using the Celltype.MSigDB gene set showed overlap with genes associated with Manno Midbrain Neurotypes HMGL, Descartes Fetal Cerebrum and Cerebellum Microglia gene sets, as well as Descartes Fetal Cerebrum and Cerebellum Vascular Endothelial Cells while suppressing expression of genes associated with Manno midbrain neurotypes HOPC, HNBGABA, HDA1/2, HRGL2A, HAD, HNBML1/5 HSERT, HRGL2b/C, HRN, HOMTN, HRGL1 and Descartes fetal cerebellum oligodendrocytes (**Figure 3E**).

### Single cell RNA sequencing

Our qRT-PCR indicated that monolayer cultures, primarily consisting of more differentiated cells, increased neuronal marker expression in response to radiation combined with forskolin treatment. However, the effect was less pronounced when performed with gliomaspheres in suspension culture which are enriched for GSCs **(Suppl. Figure 3A/B)**. We consider that the differentiated cells grown out from the surviving populations would die by anoikis under the suspension culture conditions [32]. To allow for studying GSCs and more differentiated cells in parallel we first tested if gliomaspheres grown on Poly-D-Lysine/Laminin-coated plates would still respond to irradiation combined with forskolin treatment in similar ways as mostly differentiated monolayer cells. These culture conditions maintained viable cells for 21 days irrespective of the treatment arm **(Suppl. Table 3)**. The addition of forskolin to unirradiated or irradiated cells induced neuronal marker expression (**Figure 4A, Suppl. Figure 3C/D**) and morphological changes with elongated cell bodies (**Figure 4B, black arrows).** qRT-PCR for neuronal markers revealed a significant increase in β3-tubulin expression (**Figure 4C**), and neurofilament light chain expression (**Figure 4D**). Furthermore, we observed a significant increase in the expression of the microglia marker TMEM119 (**Figure 4E**). Expression of neurofilament light chain and β3-tubulin at the protein level was confirmed by Western blotting (**Figure 4F-H)**.

**Figure 4.**
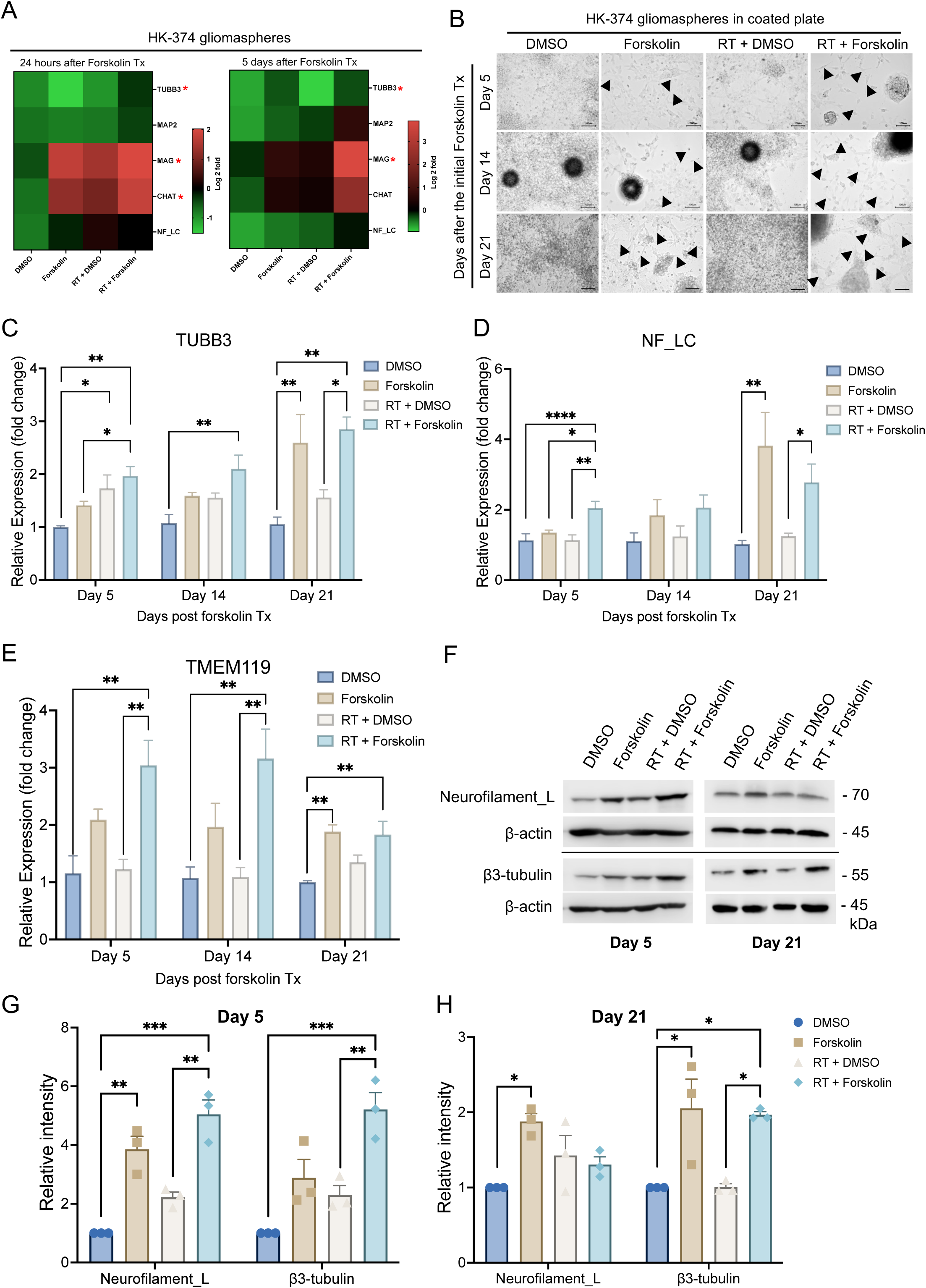
Glioma cells grown on poly-D-lysine/laminin-coated plates maintain the response to irradiation and forskolin treatment. Heatmaps showing the results of quantitative RT-PCR for the neuronal markers in HK-374 glioma spheres treated with radiation (a single dose of 4 Gy) in the presence or absence of forskolin (10 µM) for 24 hours and 5 consecutive days under suspension culture in ultra-low adhesion plates (**A**). Neuron-like cell morphology changes in HK-374 glioma spheres grown on Poly-D-Lysine/Laminin coated plates upon combine treatment of radiation and forskolin for 5, 14 and 21 days (black arrows showed the elongation of cell bodies) **(B)**. Cells could be maintained under these culture conditions for 21 days with persisting significantly increased β3-tubulin, and neurofilament light chain marker expression after irradiation and forskolin treatment (**C/D**). Additionally, we found a significant increase of the microglia marker TMEM119 after treatment with radiation and forskolin (**E**). The significant increase in neuronal marker expression was confirmed in Western blots (**F-H**). All experiments have been performed with at least 3 biological independent repeats. *p-*values were calculated using One-way ANOVA for **C-F, G, H**; The *p*-values listed in the heatmaps were from the comparison of RT + Forskolin to RT + DMSO. * *p*-value < 0.05, *\**\* *p*-value < 0.01, *** *p*-value < 0.001, **** *p*-value < 0.0001.

Our bulk RNAseq data has shown that the gene expression profile in cells treated with radiation and forskolin overlapping with a gene set associated not only with a neuronal signature but also fetal microglia and vascular endothelial cells was rather unexpected, and we therefore sought to study differences in cell fate in more details. We next performed scRNAseq using 20,028 cells treated with 0 (6,738) or 4 Gy (5,403), forskolin (5,494), or irradiation and forskolin (2,393), grown on Poly-D-Lysine/Laminin-coated plates for 5 days. Lovain clustering identified 24 unique clusters of cells that could be assigned to the different treatment arms of the study **(Suppl. Figure 4A)**. While there was some overlap between control cells (DMSO-treated) and irradiated cells and – to a lesser extent – control and forskolin-treated cells, cells treated with radiation and forskolin separated into 3 unique clusters (2, 17 and 18) **(Figure 5A)**. We next compared marker genes of the identified clusters to gene expression signatures of the adult and developing brain as described previously [24]. Correlation coefficients were generally low, and the closest normal brain cell type was assigned to individual GBM clusters **(Figure 5B)**. The main clusters in untreated control cells were primarily comprised of glycolytic-(33.4 %), inhibitory neuron- (15.3 %), NPC- (15.3 %), OPC- (13 %), and neuron-like cells (11.4 %). 5 Days after the initial forskolin treatment (7 Days after treatment with a single dose of 4 Gy) the dominant cell populations were NPC-like cells (30.2 %), inhibitory neuron-like (22.8 %) and 22.8 % of cells of low quality, the latter consistent with radiation-induced cell death. The relative increase in NPC-like cells agreed with the known radioresistant phenotype of glioma stem cells [3] and the decrease of the glycolytic-like cell population to 1.3 % was in line with our previous reports of metabolic heterogeneity and metabolic plasticity [33, 34]. Likewise, neuron-like and OPC-like cells dropped to 2 % and 3.4 %, respectively. After treatment with forskolin for five days the dominant cell clusters were vascular-like cells (37.7 %), neuron-like cells (28.3 %), astrocyte-like cells (18.1 %) and outer radial glia-like cells (oRG, 4.7 %). Combined treatment with radiation and forskolin led to the occurrence of microglia-like cells (59.1 %), neuron-like cells (22.7 %), cells in the G1/S-phase of the cell cycle (“Dividing”, 16.2 %) and a reduction in NPC-like cells (0.17 %), OPC-like cells (0.22 %) and oRG-like cells (0 %) **(Suppl. Table 4)**. Although microglia-like cells expressed immune-related genes they lacked expression of classical microglia markers **(Suppl. Figure 5).** Notably, microglia-like cells were the rarest cell population in control cells and cells treated with radiation or forskolin alone **(Figure 5C, Suppl. Table 4)**. Violin plots of marker genes for the different cell types are shown in **Figure 5D**.

**Figure 5.**
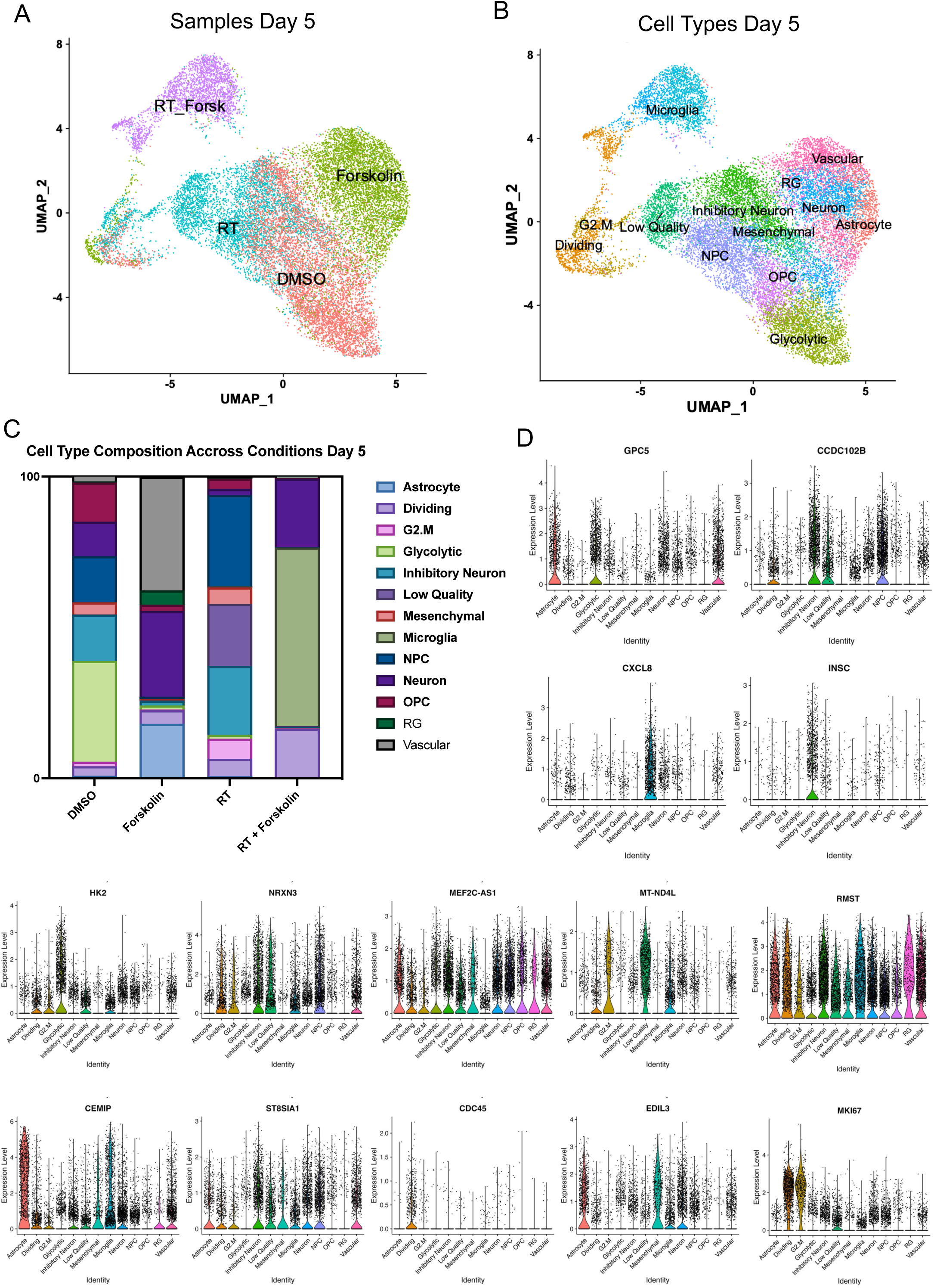
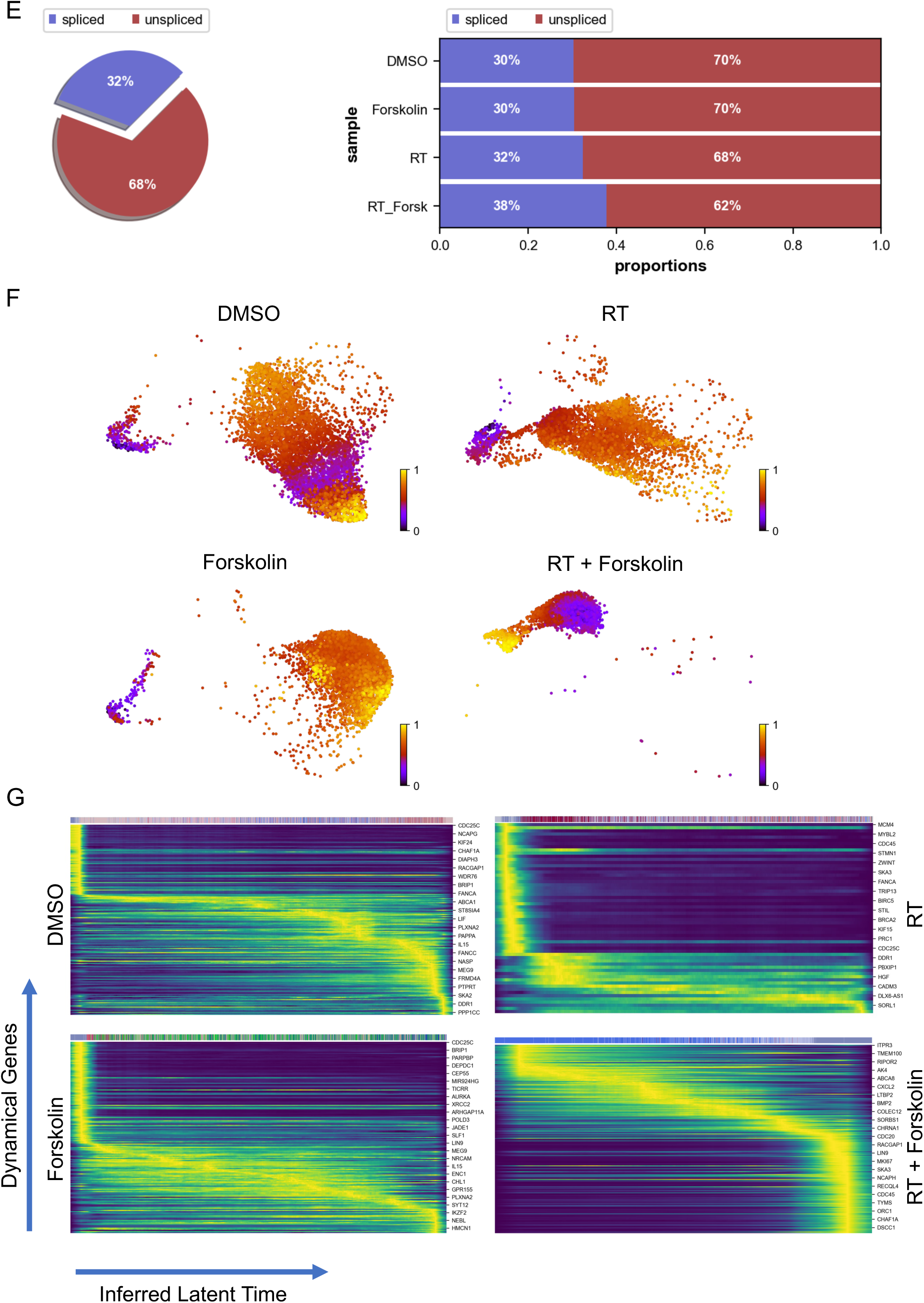
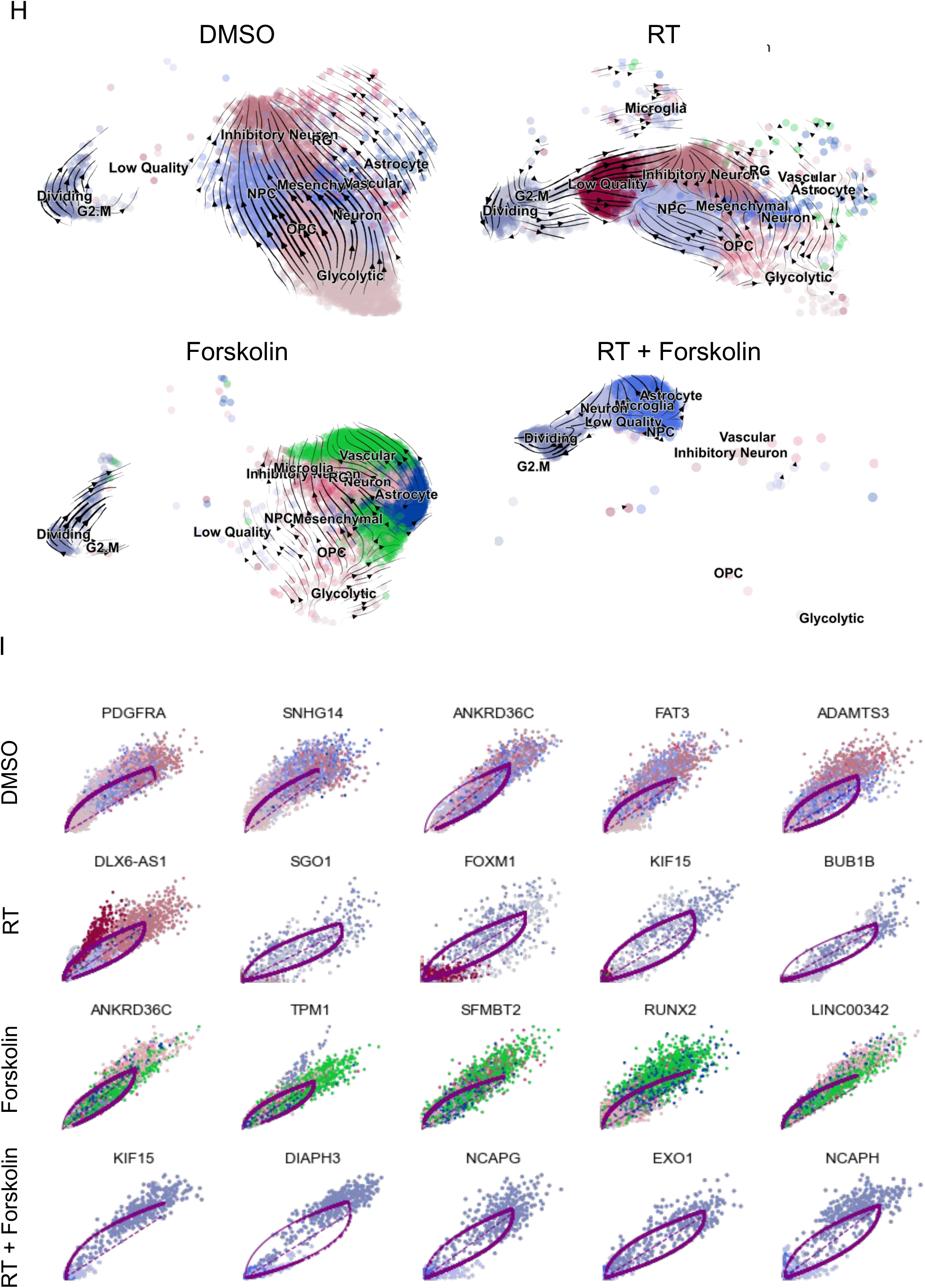
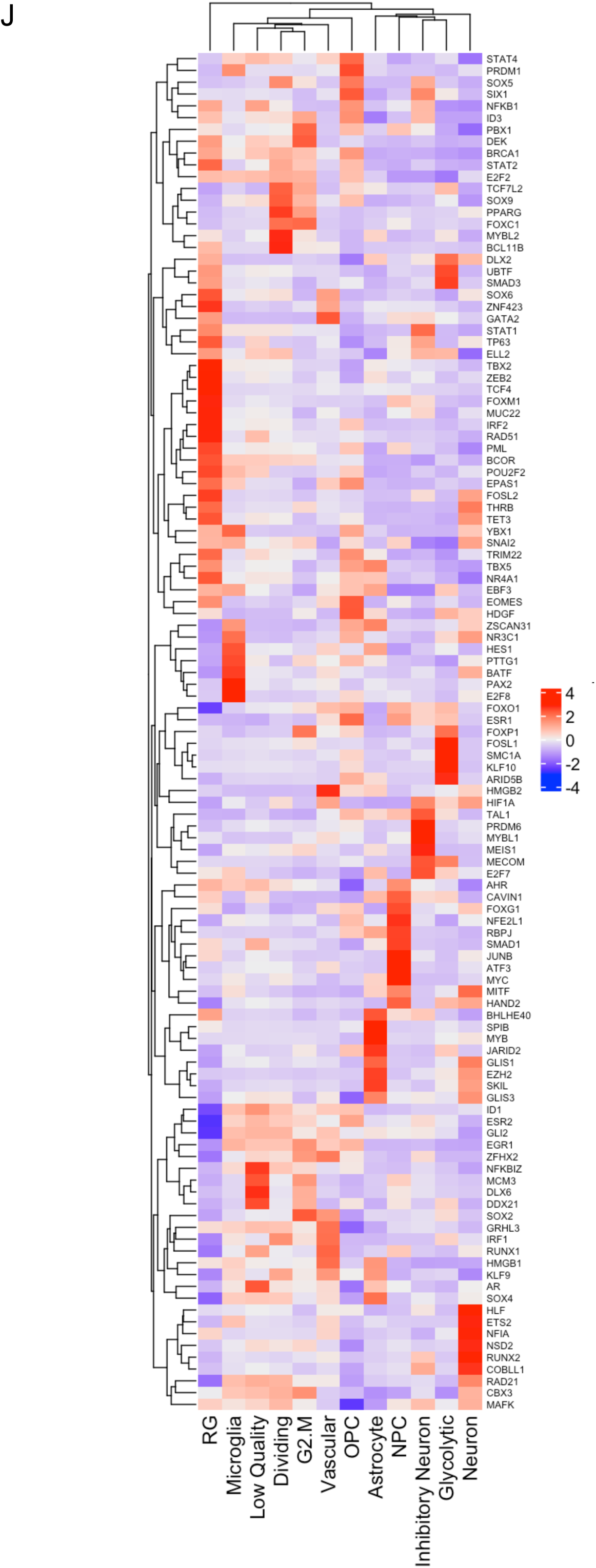
scRNAseq of HK-374 glioma cells (day 5) UMAP plots (**A**) of 24 identified clusters that could be attributed to the different treatments. Cell type annotations are shown in **B**. Stacked columns show the distribution of the different cell types for each treatment condition (**C**). Violin plots of marker genes for the identified cell types are shown in **D**. Percentage of spliced and unspliced RNA in all samples and in each treatment condition (**E**). UMAP plot of latent time for all 4 conditions (**F**). Plot of dynamical genes against inferred latent time for all samples (**G**). Trajectory analysis for all four conditions (**H**). Expression of the top 5 driver genes for each condition over latent time (**I**). Transcription factor engagement for all predicted cell types calculated using a Bayesian model (BITFAM) (**J**).

In consideration of the notable shifts in cluster composition between the different treatments we next sought to perform a trajectory analysis of the data based on RNA splicing information at day 5. Comparing the amount of spliced and unspliced RNA we found 70 % of the RNA in control cells and forskolin-treated cells unspliced, while irradiated cells showed 68 % of unspliced RNA. The amount of unspliced RNA dropped to 62 % in cells treated with radiation and forskolin **(Figure 5E)**. Next, we used scVelo’s dynamical model to compute cell trajectories based on RNA expression and splicing information by calculating RNA velocities, dynamical genes, and latent time [25] for the three different treatments and the control sample cells that were used as starting point UMAP plots of latent time suggested that control cells and cells treated with radiation or forskolin originated from dividing cells while cells treated with radiation and forskolin branched from the microglia-like cell population **(Figure 5F)**. We observed cascading dynamical genes over inferred latent time for all four conditions suggesting progression through differentiation of dedifferentiation steps **(Figure 5G)**, which was supported by the trajectory analysis of the cells **(Figure 5H)**. The trajectory analysis indicated dynamic conversion of cell types in untreated control cells consistent with the known intratumoral cellular heterogeneity of GBM **(Figure 5H)**. As expected, cells in the control sample developed into the different phenotypes from the dividing cell population. The main trajectory led to glycolytic cells from which a subset further developed into OPC-like, neuron-like and NPC-like cells and into inhibitory neuron-like cells. This trajectory was maintained in irradiated cells, except for the occurrence of ‘low quality’ cells originating directly from the dividing cell population **(Figure 5H)**. Treatment of the cells with forskolin caused a deviation from this trajectory with dividing cells transitioning into vascular-like cells, neuron-like cells, astrocyte-like cells, and outer radial glia-like cells **(Figure 5H)**.

Lastly, treatment with radiation and forskolin completely altered the trajectory of the cells with the appearance of microglia-like cells and dividing cells leading into neuron-like cells, thereby omitting all other phenotypes **(Figure 5H)**. The top 5 driver genes for each condition are shown in **Figure 5I**.

Next we sought to identify transcription factors engaged in changes of gene expression profiles in individual cell types using a Bayesian model [26] **(Figure 5J)**. For microglia-like cells we calculated engagement of HAND2, MITF, MYC, ATF3, JUNB, SMAD1, RBPJ, NFE2L1, FOXG1, CAVIN1, AHR, TAL1, ESR1, FOXO1, SNAI2, PML, FOXM1, GATA2 and PBX1. None of these transcription factors are defining microglia cells but rather pointed to proinflammatory signaling (MYC, ATF3, JUNB, SMAD1, FOXO1, AHR, ESR1) [35–41].

For neuron-like cells we calculated engagement of MAFK, CBX3, Rad21, COBLL1, RUNX2, NSD2, NFIA, ETS2, HLF, GLIS3, SKIL, EZH2, GLIS1, HAND2, MITF, FOXG1, HIF1A, HMGB2, BATF, NR3C1, ZSCAN31, HDGF, SNAI2, YBX1, TET3, THRB, FOSL2 and. EZH2 [42] and FOSL2 [43] are associated with synaptic plasticity and FOXG1 [44], DLX2 [45], THRB [46] participate in CNS development.

Finally, we studied how stable these phenotypes would be over time. Repeating the scRNAseq experiment with cells treated for 21 days (25 unique clusters of cells were identified by Lovain clustering, **Suppl. Figure 4B**) and projecting cell types identified in day 5 samples onto clusters from day 21 **(Figure 6A)**, we observed that the culture conditions changed the overall composition of control cells with mesenchymal-like cells (25.8 %), dividing cells (19.3 %), NPC-like cells (13.4 %), astrocyte-like cells (12.7 %), or neuron-like cells (10.6 %) as the dominant cell types and oRG-like and OPC-like cells no longer present in any of the treatment arms. At day 21, irradiated cells had largely redistributed to match the cluster distribution of control cells but still maintained an increase in NPC-like cells (19.8 %). Forskolin-treated cells predominantly showed and astrocyte-like (38.2 %), neuron-like (25.3 %), mesenchymal-like (11.4 %), or NPC-like (10.7 %) phenotype. The dominant phenotypes in cells treated with radiation plus forskolin remained microglia-like cells (39 %), followed by NPC-like (15.5 %), and neuron-like cells (10.2 %) **(Figure 6B/C; Suppl. Table 5)**.

**Figure 6.**
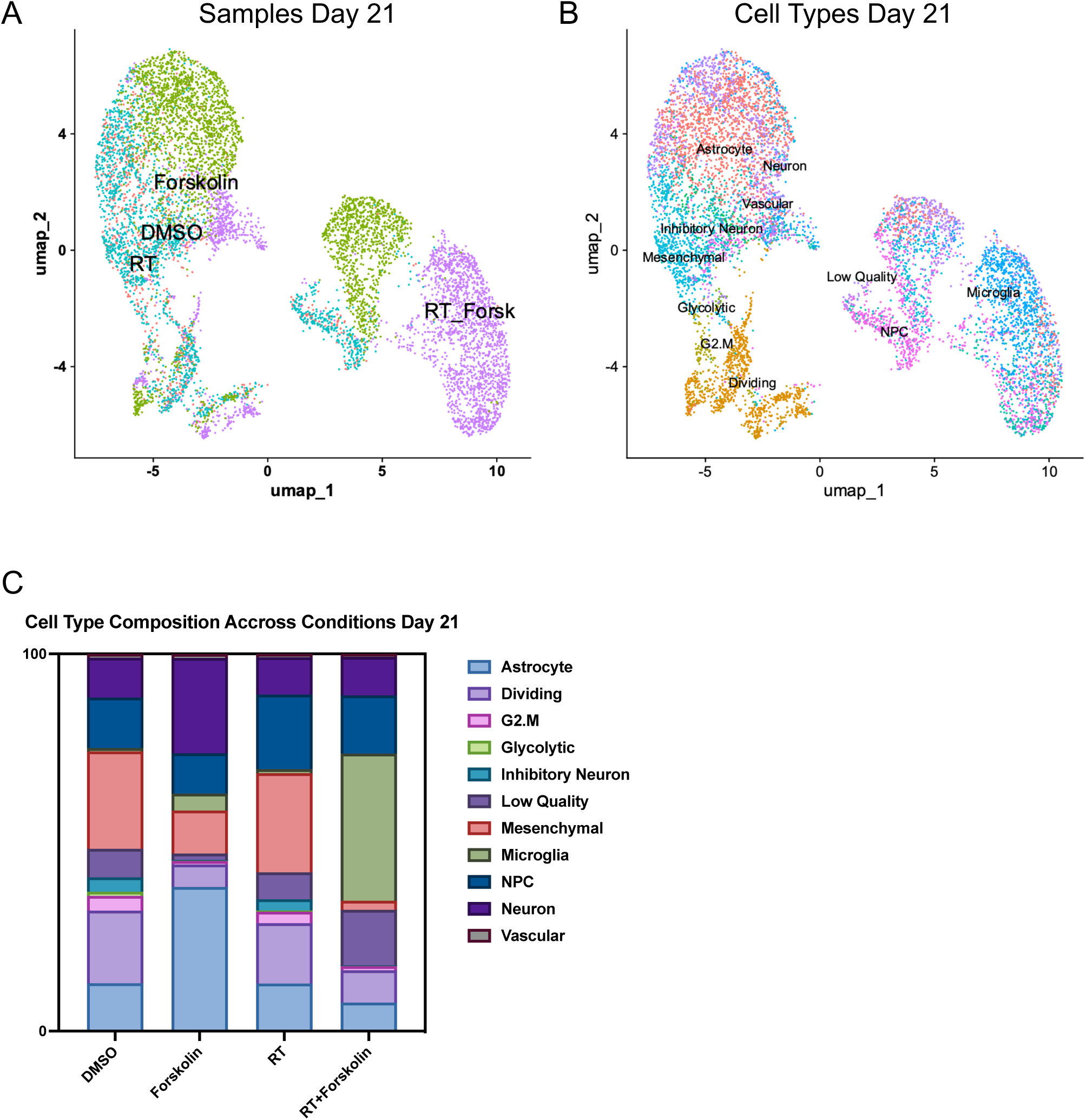
scRNAseq of HK-374 glioma cells (day 21) UMAP plot of clusters identified 21 days after start of forskolin treatment attributed to the four different conditions (**A**) and projection of cell types identified on day 5 (**B**). Stacked columns show the distribution of the different cell types for each treatment condition (**C**).

### A combination of radiation and forskolin improves median survival in vivo

*In silico*, forskolin has an estimated LogBB of -0.24 [47], indicating its ability to cross the blood-brain barrier [48]. Therefore, we tested if the combination of radiation and forskolin would affect the number of GSCs *in vivo*. HK374 cells were implanted into the striatum of NSG mice. After 3 days of grafting, tumors were irradiated with 0 of 4 Gy. After 48 hours, the animals were treated with daily forskolin injections (5 mg/kg) on a 5-day on/2-day off schedule for 2 weeks. Tumors were harvested, digested into single cell suspensions, and subjected to clonal sphere-formation assays for 10 days **(Figure 7A)**. The combination of 4 Gy and Forskolin led to a significant reduction in sphere-forming capacity from 5.25 % to 0.24 % (*p*=0.015). Treatment with radiation alone also significantly reduced sphere-formation, while forskolin alone had no significant effect (**Figure 7B/C**). Next, we calculated GSC frequencies using an ELDA. Both irradiation and forskolin treatment significantly reduced the number of GSCs, while the combination of radiation and forskolin significantly reduced GSC numbers to 0.04 %, 95 % class interval: 0.03 % to 0.05 %, *p*<0.0001 **(Figure 7D/E, Suppl. Tables 6/7)**. Confocal imaging of corresponding tumor sections revealed that forskolin treatment and combined treatment with radiation and forskolin induced the expression of neurofilament light chain and Tuj1 in GFP-expressing GBM cells *in vivo* **(Figure 7F, white arrows)**.

**Figure 7.**
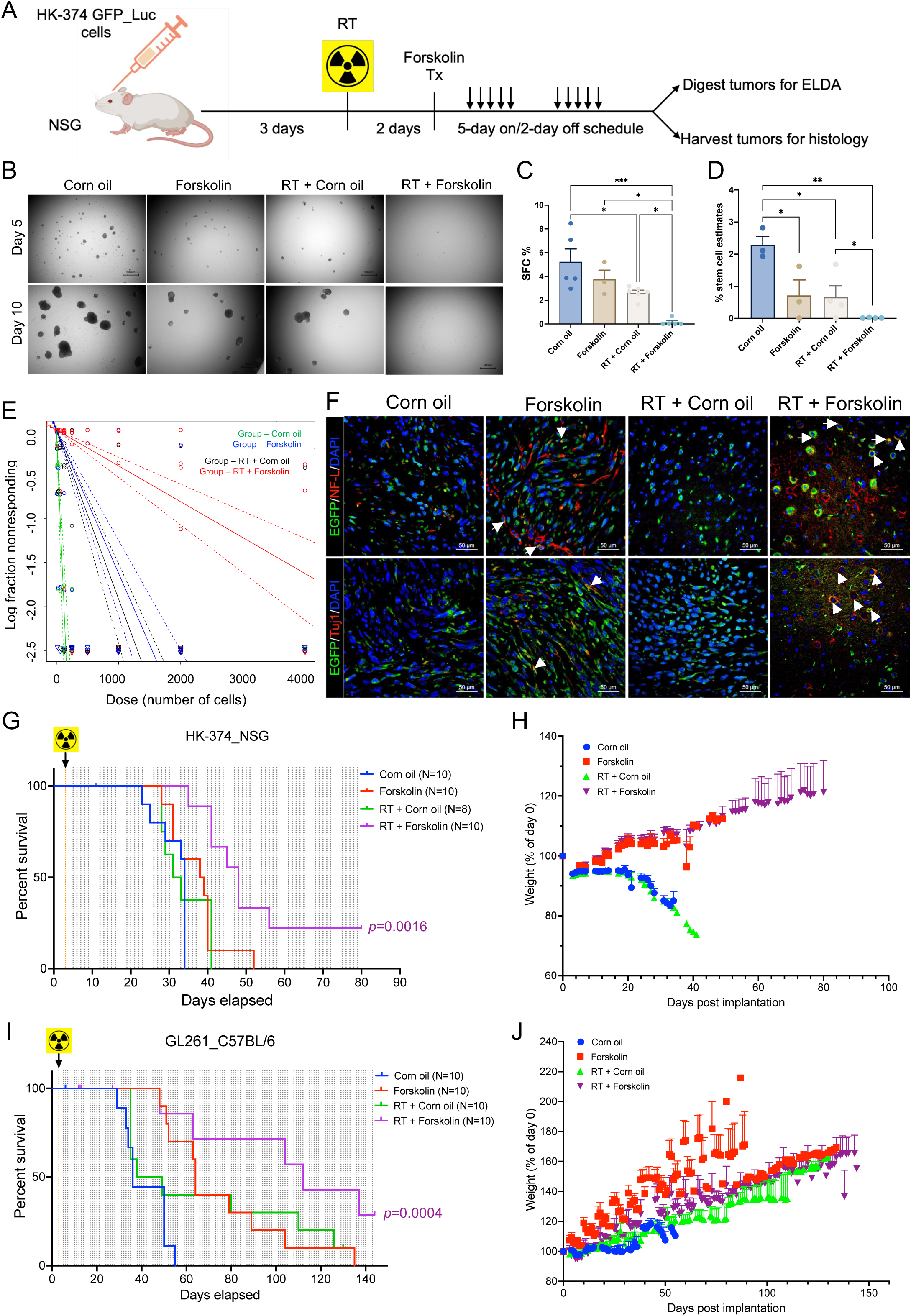
A combination of radiation and forskolin eliminates GSC *in vivo* and prolongs median survival in mouse models of glioblastoma. Schematic of the experimental design for Figure 7 (**A**). Single cell suspension from the tumors harvested showed a significant reduction in sphere-formation for animals treated with radiation, while a combination of radiation and forskolin almost completely prevented sphere-formation (**B/C**). Using an ELDA we found that forskolin treatment and irradiation alone significantly reduced the frequency of GSCs, while the combined treatment with radiation and forskolin nearly eliminated all GSCs (**D/E**). Confocal imaging of tumor section of the animals revealed expression of neurofilament light chain and Tuj-1 in a subset of GFP-expressing tumor cells (**F**, white arrows: colocalization of double stained cells). Treatment of HK-374 PDOXs in NSG mice (**G/H**) and intracranial syngeneic GL261 tumors in C57Bl/6 mice (**I/J**) with a single dose of 10 Gy and forskolin (5 mg/kg) was well tolerated and significantly prolonged the median survival of the animals. All experiments have been performed with at least 3 biological independent repeats. *p-*values were calculated using One-way ANOVA for **C**; *p-*values were calculated and generated by ELDA software for **D**; Log-rank (Mantel-Cox) test for comparison of Kaplan-Meier survival curves in **G, I.** * *p*-value < 0.05, *\**\* *p*-value < 0.01, *** *p*-value < 0.001.

Finally, we tested if the combination of radiation and forskolin would lead to an improved median survival in mouse models of glioma. HK374 cells were implanted into the striatum of NSG mice. After 3 days of grafting, PDOXs were irradiated with a single dose of 10 Gy, equivalent to a total dose of 18 Gy in 2 Gy fractions. Control mice were sham irradiated. Forty-eight hours later, mice were treated with forskolin at 5 mg/kg (5 days on/2 days off schedule) until they reached euthanasia criteria. Control animals were treated with solvent only. While radiation or forskolin treatment alone had no effect in this highly aggressive and rapidly growing PDOX, the combination of radiation and forskolin significantly increased the median survival of the animals from 34 to 48 days (**Figure 7G**; *p*<0.0001, Log-Rank test). Forskolin treatment was well tolerated, and animals continued to gain weight after combined treatment **(Figure 7H)**.

In the syngeneic GL261 glioma model in C57Bl/6 mice control mice had a median survival of 36 days. Median survival was increased to 43.5 days in irradiated mice (not significant), to 64 days in mice treated with forskolin (*p*=0.001, Log-Rank test) and to 129 days in mice receiving the combination of radiation and forskolin (**Figure 7I**; *p*=0.0021, Log-Rank test). As in NSG mice the addition of forskolin was well tolerated **(Figure 7J)**.

## Discussion

In our current study we hypothesized that glioma cells, that survive irradiation, go through a transient state of multipotency that can be exploited to drive GBM cells into a postmitotic neuron-like state. Our hypothesis was based on our previous observations that differentiated non-stem glioma cells that survive exposure to ionizing radiation respond with a phenotype conversion into induced GSCs [15] while surviving, preexisting GSCs transdifferentiated into endothelial- and pericyte-like cells [18]. The underlying mechanisms were global epigenetic remodeling through changes in histone methylation and acetylation and subsequent changes in open chromatin [15, 18], with the latter mediated by the histone acetyltransferase EP300 [18]. Induced GSCs and transdifferentiated pericyte- and endothelial-like cells contributed to tumor recurrences and treatment resistance [15, 18] and preventing phenotype conversion or transdifferentiation prolonged median survival in PDOXs mouse models of GBM [15, 18, 49, 50]. We show here that the addition of forskolin to radiation drives GBM cells into microglia- and neuron-like states, reduced cell proliferation and the number of functional GSCs *in vitro* and *in vivo* and prolonged the median survival in syngeneic and PDOX mouse models of GBM.

Several previous studies have attempted differentiation therapies for GBM. Most studies succeeded to some extent *in vitro* and cells showed decreased tumorigenicity when implanted into mice. For example, all-trans retinoic acid (ATRA) successfully differentiated GSCs *in vitro* and reduced their tumorigenicity *in vivo* [51] but clinical translation targeting GBM with ATRA has consistently failed [52, 53]. Another study induced GSC differentiation through stereotactical injection of a WNT inhibitor, an SHH inhibitor and BMP [54]. While effective, the intracranial application route could be clinically challenging and is likely to miss GBM cells, dispersed into the normal parenchyma that are not in reach of the injections.

Forskolin and cAMP have long been known to induce glioma differentiation *in vitro* [55, 56] and the use of forskolin or dbcAMP to induce terminal neuronal differentiation of glioma *in vivo* has been previously reported [57]. Like in our current study, this treatment alone had minimal effects on median survival even when combined with a Wnt inhibitor and even though dbcAMP was given at a high dose [57]. Our scRNAseq data indicate that forskolin treatment alone not only induced neuron-, astrocyte-, and vascular-like cells but also a small number of cells with an outer radial glia-like cell state (4.7 %), a cell population present in small numbers in control and irradiated cells but completely absent in cells treated with radiation and forskolin. It was previously reported that oRG cells have glioma stem cells features [24] and this observation could explain why dbcAMP or forskolin treatment alone have limited impact on median survival in glioma models.

Radiation therapy is and -for the foreseeable time- will remain the most effective treatment modality against GBM and any novel treatment modality will most like be added to standard of care, surgery, and radiation, or will at least benchmarked against it. Our discovery, that radiation induces a transient multipotent state added a previously unknown facete to the radiation response of cancer, that leads to increased cellular plasticity, which is increasingly recognized as an emerging hallmark of cancer [58]. This increase in plasticity leads to dedifferentiation or transdifferentiation and is by itself detrimental with respect to tumor control. By exploiting radiation-induced multipotency our approach highjacks a unique feature of surviving cells that is inevitably induced by a main pillar of the current standard of care. With the addition of forskolin to radiation treatment we utilize an established method that forces iPS cells into neuronal differentiation.

However, GBM cells, that gain a certain level of multipotency, and iPS cells do not share the same level of multipotency, drastically differ in mutational burden and irradiated GBM cells continue to signal through the DNA damage response. Hence, it is not expected that forced differentiation of GBM cell yields in *bona fide* induced neurons. Our finding that the combination of radiation and forskolin predominately led to the occurrence of microglia-like cells and to a lesser extent to neuron-like cells was unexpected. The role of this predominant immune cell-like phenotype in the presence of an intact immune system needs further investigation. However, a combination of radiation and forskolin was superior to their individual effects on inducing neuronal marker expression and inhibiting cell proliferation of GBM cells, reducing self-renewal capacity of GSCs and GSC frequency and prolonging median survival of glioma-bearing mice and leading to long term tumor control in some of the mice. This suggested that this combination therapy induced cell phenotype that can no longer sustain tumor growth. The very low dose of forskolin used here was well tolerated and the human equivalent dose (25 mg for 60 kg body weight) is far below the usual supplement dose of 50mg/day for forskolin. With forskolin crossing the blood-brain barrier and its long use in ayurvedic medicine, this combination therapy can easily be translated into a clinical trial.

## Acknowledgements

This work used computational and storage services associated with the Hoffman2 Shared Cluster provided by UCLA Office of Advanced Research Computing’s Research Technology Group.

This work was made possible, in part, through access to the following: the Genomics Research and Technology Hub (formerly Genomics High-Throughput Facility) Shared Resource of the Cancer Center Support Grant (P30CA-062203), the Single Cell Analysis Core shared resource of Complexity, Cooperation and Community in Cancer (U54CA217378), the Genomics-Bioinformatics Core of the Skin Biology Resource Based Center @ UCI (P30AR075047) at the University of California, Irvine and NIH shared instrumentation grants 1S10RR025496-01, 1S10OD010794-01, and 1S10OD021718-01.

## Notes

Funding: FP was supported by grants from the National Cancer Institute (R01CA260886, R01CA281682) and the UCLA Eli and Edythe Broad Center of Regenerative Medicine and Stem Cell Research Award Program. FP and HIK were supported by the California Institute for Regenerative Medicine (CIRM; DISC2-14083).

AB, HIK and FP were supported by the American Cancer Society (<u>CSCC-Team-23-980262-01-CSCC</u>).

### Competing Interest Statement

The authors have declared no competing interest.

## References

1. Hemmati, H.D., et al., Cancerous stem cells can arise from pediatric brain tumors. Proc Natl Acad Sci U S A, 2003. 100(25): p. 15178–83.

2. Singh, S.K., et al., Identification of a cancer stem cell in human brain tumors. Cancer Res, 2003. 63(18): p. 5821–8.

3. Bao, S., Glioma stem cells promote radioresistance by preferential activation of the DNA damage response. Nature, 2006. 444: p. 756–760.

4. Eramo, A., et al., Chemotherapy resistance of glioblastoma stem cells. Cell Death Differ, 2006. 13(7): p. 1238–41.

5. Yao, M., et al., Cellular origin of glioblastoma and its implication in precision therapy. Cell Mol Immunol, 2018. 15(8): p. 737–739.

6. Swartling, F.J., et al., Deregulated proliferation and differentiation in brain tumors. Cell Tissue Res, 2015. 359(1): p. 225–54.

7. Yan, K., et al., Glioma cancer stem cells secrete Gremlin1 to promote their maintenance within the tumor hierarchy. Genes Dev, 2014. 28(10): p. 1085–100.

8. Yan, M., et al., IKKalpha restoration via EZH2 suppression induces nasopharyngeal carcinoma differentiation. Nat Commun, 2014. 5: p. 3661.

9. Munster, P.N., et al., The histone deacetylase inhibitor suberoylanilide hydroxamic acid induces differentiation of human breast cancer cells. Cancer Res, 2001. 61(23): p. 8492–7.

10. Basu-Roy, U., et al., PPARgamma agonists promote differentiation of cancer stem cells by restraining YAP transcriptional activity. Oncotarget, 2016. 7(38): p. 60954–60970.

11. Ishay-Ronen, D., et al., Gain Fat-Lose Metastasis: Converting Invasive Breast Cancer Cells into Adipocytes Inhibits Cancer Metastasis. Cancer Cell, 2019. 35(1): p. 17–32 e6.

12. Storm, E.E., et al., Targeting PTPRK-RSPO3 colon tumours promotes differentiation and loss of stem-cell function. Nature, 2016. 529(7584): p. 97–100.

13. Ishay-Ronen, D. and G. Christofori, Targeting Cancer Cell Metastasis by Converting Cancer Cells into Fat. Cancer Res, 2019. 79(21): p. 5471–5475.

14. Lagadec, C., et al., Radiation-induced reprogramming of breast cancer cells. Stem Cells, 2012. 30(5): p. 833–44.

15. Bhat, K., et al., The dopamine receptor antagonist trifluoperazine prevents phenotype conversion and improves survival in mouse models of glioblastoma. Proc Natl Acad Sci U S A, 2020. 117(20): p. 11085–11096.

16. Pisco, A.O. and S. Huang, Non-genetic cancer cell plasticity and therapy-induced stemness in tumour relapse: ’What does not kill me strengthens me’. Br J Cancer, 2015. 112(11): p. 1725–32.

17. Chen, X., et al., Induced cancer stem cells generated by radiochemotherapy and their therapeutic implications. Oncotarget, 2017. 8(10): p. 17301–17312.

18. Muthukrishnan, S.D., et al., P300 promotes tumor recurrence by regulating radiation-induced conversion of glioma stem cells to vascular-like cells. Nat Commun, 2022. 13(1): p. 6202.

19. Zhang, X., et al., Terminal differentiation and loss of tumorigenicity of human cancers via pluripotency-based reprogramming. Oncogene, 2013. 32(18): p. 2249–60, 2260 e1-21.

20. Laks, D.R., et al., Large-scale assessment of the gliomasphere model system. Neuro Oncol, 2016. 18(10): p. 1367–78.

21. Vlashi, E., et al., In vivo imaging, tracking, and targeting of cancer stem cells. Journal of the National Cancer Institute, 2009. 101(5): p. 350–9.

22. Hu, Y. and G.K. Smyth, ELDA: extreme limiting dilution analysis for comparing depleted and enriched populations in stem cell and other assays. J Immunol Methods, 2009. 347(1-2): p. 70–8.

23. Ge, S.X., E.W. Son, and R. Yao, iDEP: an integrated web application for differential expression and pathway analysis of RNA-Seq data. BMC Bioinformatics, 2018. 19(1): p. 534.

24. Bhaduri, A., et al., Outer Radial Glia-like Cancer Stem Cells Contribute to Heterogeneity of Glioblastoma. Cell Stem Cell, 2020. 26(1): p. 48–63 e6.

25. Bergen, V., et al., Generalizing RNA velocity to transient cell states through dynamical modeling. Nat Biotechnol, 2020. 38(12): p. 1408–1414.

26. Gao, S., Y. Dai, and J. Rehman, A Bayesian inference transcription factor activity model for the analysis of single-cell transcriptomes. Genome Res, 2021. 31(7): p. 1296–1311.

27. Li, W., et al., Rapid induction and long-term self-renewal of primitive neural precursors from human embryonic stem cells by small molecule inhibitors. Proc Natl Acad Sci U S A, 2011. 108(20): p. 8299–304.

28. Chambers, S.M., et al., Highly efficient neural conversion of human ES and iPS cells by dual inhibition of SMAD signaling. Nat Biotechnol, 2009. 27(3): p. 275–80.

29. Seamon, K.B. and J.W. Daly, Forskolin: its biological and chemical properties. Adv Cyclic Nucleotide Protein Phosphorylation Res, 1986. 20: p. 1–150.

30. Muraguchi, A., et al., Inhibition of human B cell activation by diterpine forskolin: interference with B cell growth factor-induced G1 to S transition of the B cell cycle. J Immunol, 1984. 133(3): p. 1283–7.

31. McBride, W.H., et al., NF-kappa B, cytokines, proteasomes, and low-dose radiation exposure. Mil Med, 2002. 167(2 Suppl): p. 66–7.

32. Ray, S., Tumorsphere Formation Assay: A Cancer Stem-Like Cell Characterization in Pediatric Brain Cancer Medulloblastoma. Methods Mol Biol, 2023. 2701: p. 253–259.

33. Vlashi, E., et al., Metabolic state of glioma stem cells and nontumorigenic cells. Proceedings of the National Academy of Sciences of the United States of America, 2011. 108(38): p. 16062–16067.

34. Vlashi, E., et al., Metabolic differences in breast cancer stem cells and differentiated progeny. Breast Cancer Res Treat, 2014. 146(3): p. 525–34.

35. Gao, L., et al., Baicalein Attenuates Neuroinflammation in LPS-Treated BV-2 Cells by Inhibiting Glycolysis via STAT3/c-Myc Pathway. Neurochem Res, 2023. 48(11): p. 3363–3377.

36. Zhang, Y., et al., Microglia-specific transcriptional repression of interferon-regulated genes after prolonged stress in mice. Neurobiol Stress, 2022. 21: p. 100495.

37. Woo, M.S., et al., Selective modulation of lipopolysaccharide-stimulated cytokine expression and mitogen-activated protein kinase pathways by dibutyryl-cAMP in BV2 microglial cells. Brain Res Mol Brain Res, 2003. 113(1-2): p. 86–96.

38. Huang, W., et al., TGF-beta1/SMADs signaling involved in alleviating inflammation induced by nanoparticulate titanium dioxide in BV2 cells. Toxicol In Vitro, 2022. 80: p. 105303.

39. Li, Z., et al., Foxo1-mediated inflammatory response after cerebral hemorrhage in rats. Neurosci Lett, 2016. 629: p. 131–136.

40. Quintana, F.J., Regulation of central nervous system autoimmunity by the aryl hydrocarbon receptor. Semin Immunopathol, 2013. 35(6): p. 627–35.

41. Sarvari, M., et al., Estrogens regulate neuroinflammatory genes via estrogen receptors alpha and beta in the frontal cortex of middle-aged female rats. J Neuroinflammation, 2011. 8: p. 82.

42. Zhang, M., et al., Neuronal Histone Methyltransferase EZH2 Regulates Neuronal Morphogenesis, Synaptic Plasticity, and Cognitive Behavior in Mice. Neurosci Bull, 2023. 39(10): p. 1512–1532.

43. Levran, O., et al., Synaptic Plasticity and Signal Transduction Gene Polymorphisms and Vulnerability to Drug Addictions in Populations of European or African Ancestry. CNS Neurosci Ther, 2015. 21(11): p. 898–904.

44. Hettige, N.C. and C. Ernst, FOXG1 Dose in Brain Development. Front Pediatr, 2019. 7: p. 482.

45. Bulfone, A., et al., Spatially restricted expression of Dlx-1, Dlx-2 (Tes-1), Gbx-2, and Wnt-3 in the embryonic day 12.5 mouse forebrain defines potential transverse and longitudinal segmental boundaries. J Neurosci, 1993. 13(7): p. 3155–72.

46. Jones, I., et al., The thyroid hormone receptor beta gene: structure and functions in the brain and sensory systems. Thyroid, 2003. 13(11): p. 1057–68.

47. Shaker, B., et al., A machine learning-based quantitative model (LogBB_Pred) to predict the blood-brain barrier permeability (logBB value) of drug compounds. Bioinformatics, 2023. 39(10).

48. Dyabina, A.S., et al., Prediction of blood-brain barrier permeability of organic compounds. Dokl Biochem Biophys, 2016. 470(1): p. 371–374.

49. Bhat, K., et al., Dopamine Receptor Antagonists, Radiation, and Cholesterol Biosynthesis in Mouse Models of Glioblastoma. J Natl Cancer Inst, 2021. 113(8): p. 1094–1104.

50. He, L., et al., Effects of the DRD2/3 antagonist ONC201 and radiation in glioblastoma. Radiother Oncol, 2021. 161: p. 140–147.

51. Campos, B., et al., Differentiation therapy exerts antitumor effects on stem-like glioma cells. Clin Cancer Res, 2010. 16(10): p. 2715–28.

52. Kaba, S.E., et al., The treatment of recurrent cerebral gliomas with all-trans-retinoic acid (tretinoin). J Neurooncol, 1997. 34(2): p. 145–51.

53. Phuphanich, S., et al., All-trans-retinoic acid: a phase II Radiation Therapy Oncology Group study (RTOG 91-13) in patients with recurrent malignant astrocytoma. J Neurooncol, 1997. 34(2): p. 193–200.

54. Jin, X., et al., Inhibition of ID1-BMPR2 Intrinsic Signaling Sensitizes Glioma Stem Cells to Differentiation Therapy. Clin Cancer Res, 2018. 24(2): p. 383–394.

55. Chen, T.C., et al., Up-regulation of the cAMP/PKA pathway inhibits proliferation, induces differentiation, and leads to apoptosis in malignant gliomas. Lab Invest, 1998. 78(2): p. 165–74.

56. Sharma, S.K. and A.B. Raj, Transient increase in intracellular concentration of adenosine 3’:5’-cyclic monophosphate results in morphological and biochemical differentiation of C6 glioma cells in culture. J Neurosci Res, 1987. 17(2): p. 135–41.

57. Chen, Z., et al., Disruption of beta-catenin-mediated negative feedback reinforces cAMP-induced neuronal differentiation in glioma stem cells. Cell Death Dis, 2022. 13(5): p. 493.

58. Hanahan, D., Hallmarks of Cancer: New Dimensions. Cancer Discov, 2022. 12(1): p. 31–46.

