## Supplemental Materials for "Radiation-Induced Cellular Plasticity: A Strategy for Combatting Glioblastoma"

**Table of contents:**

**1. Supplementary Figures**

**2. Supplementary Tables**

**Supplementary Figures**

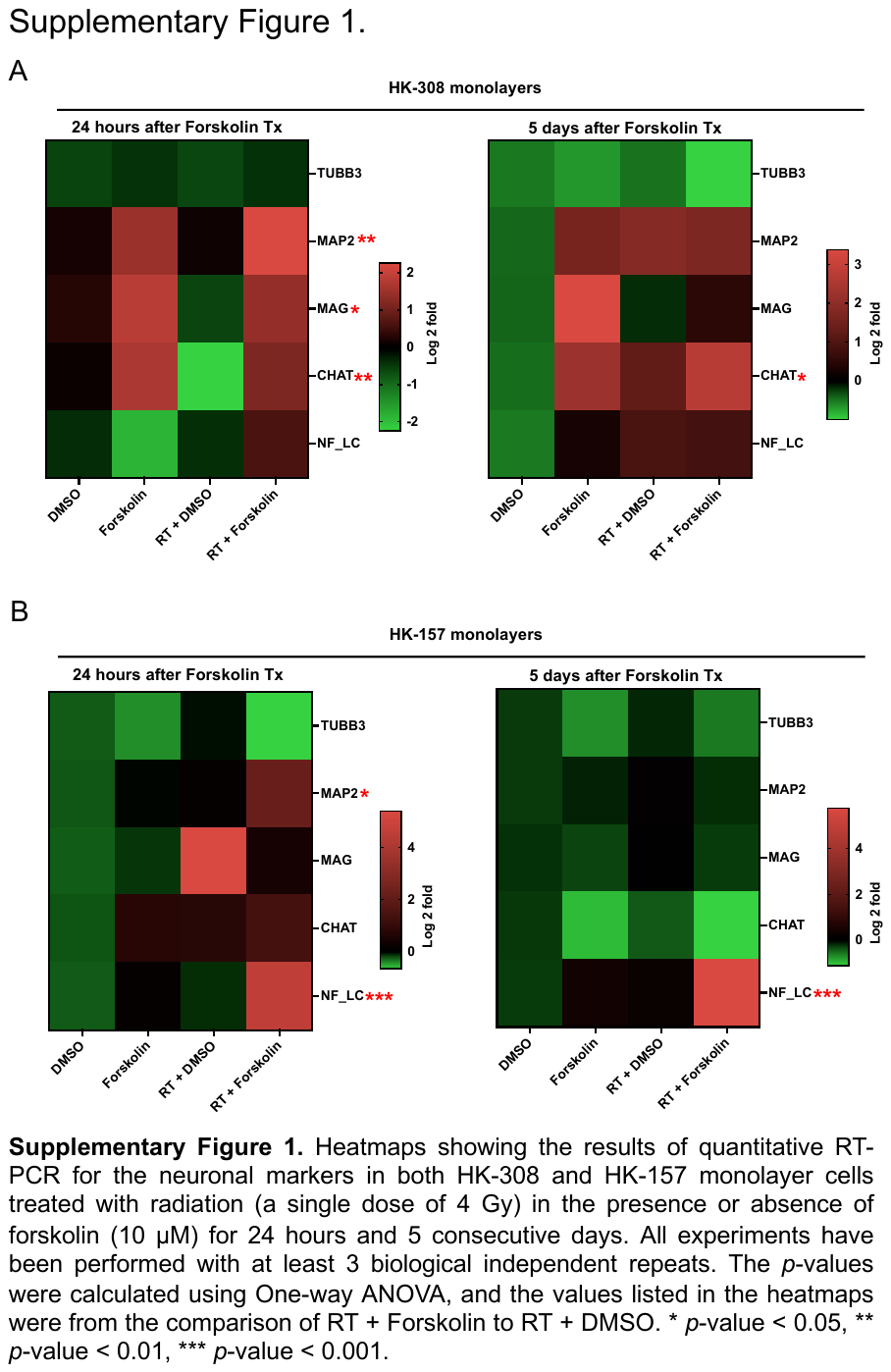

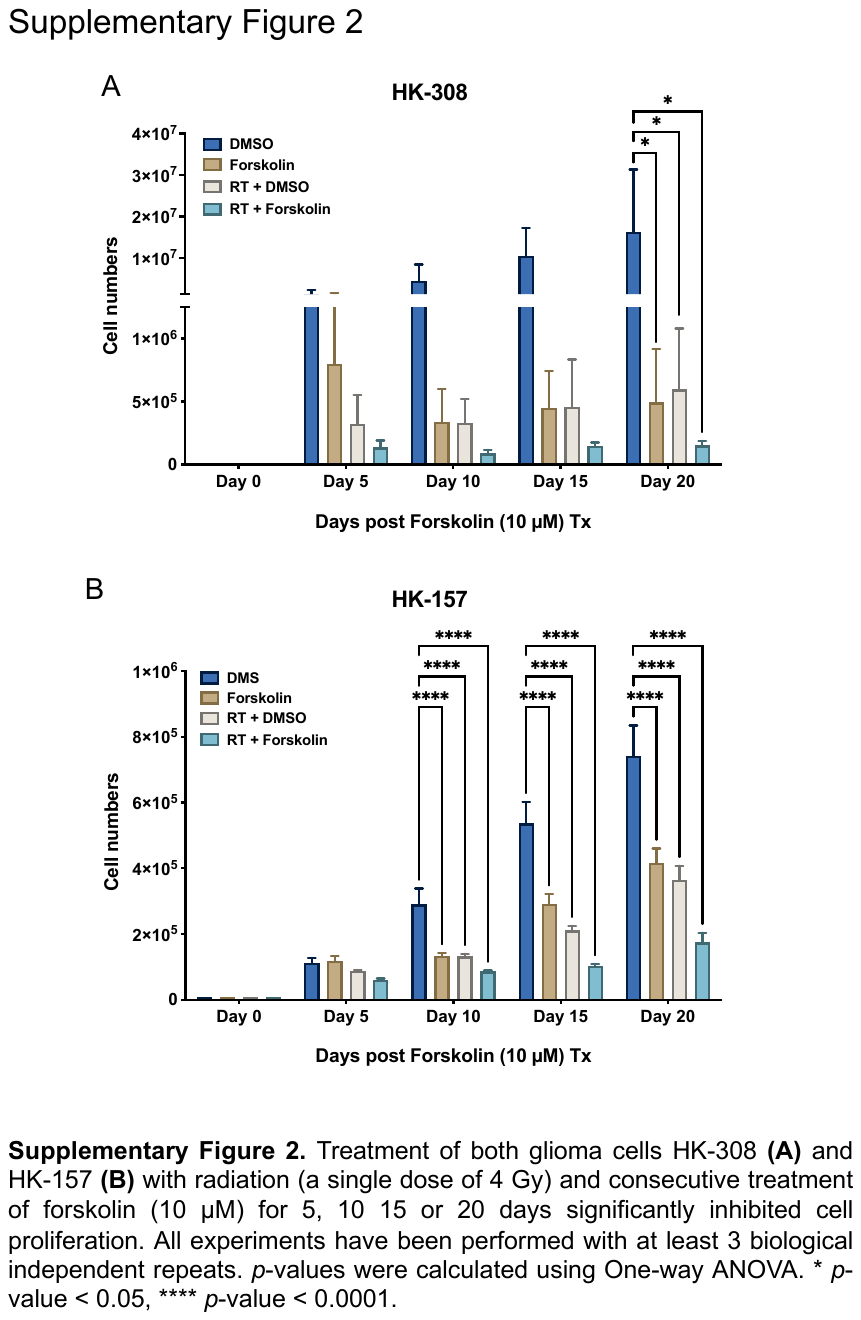

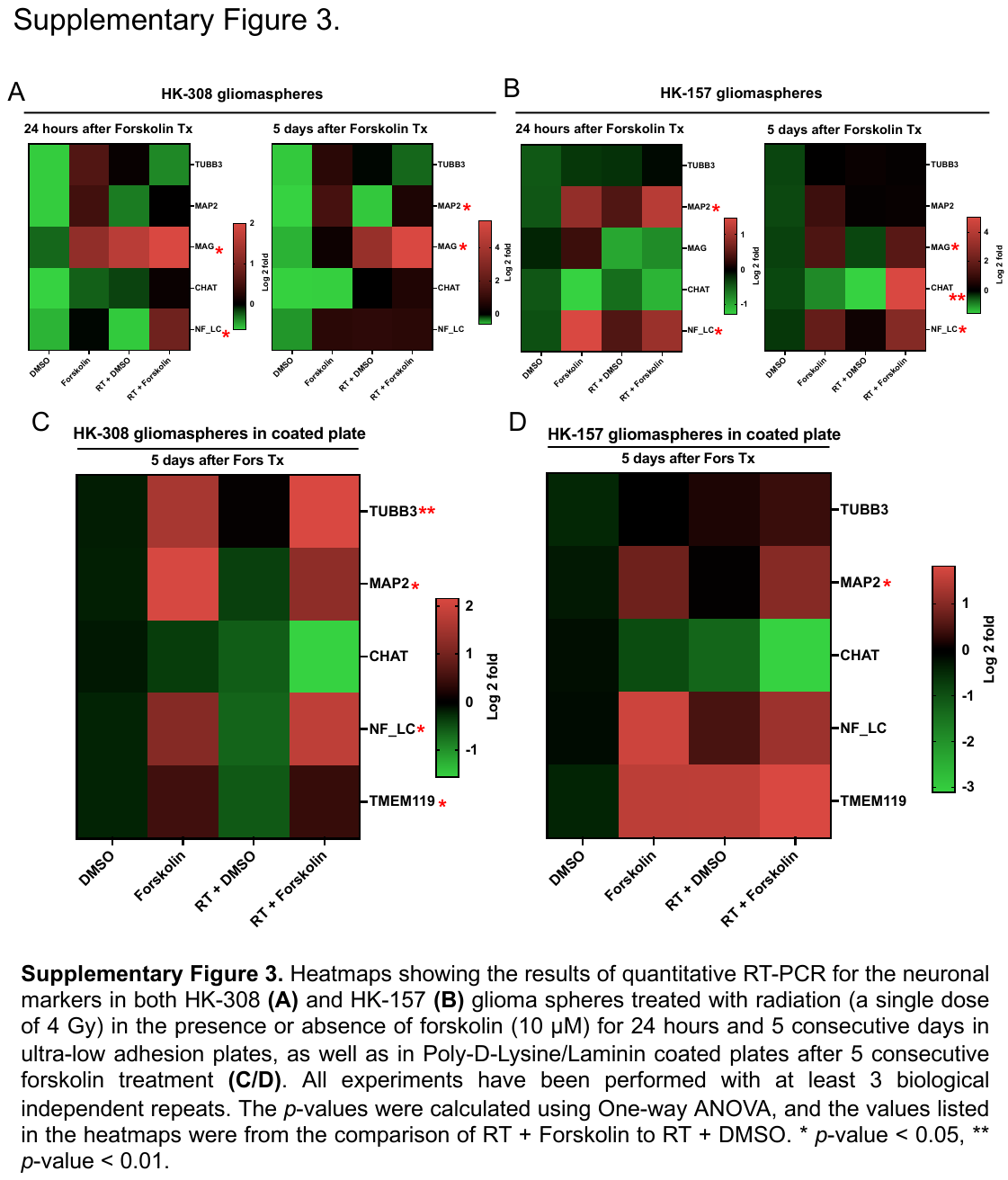

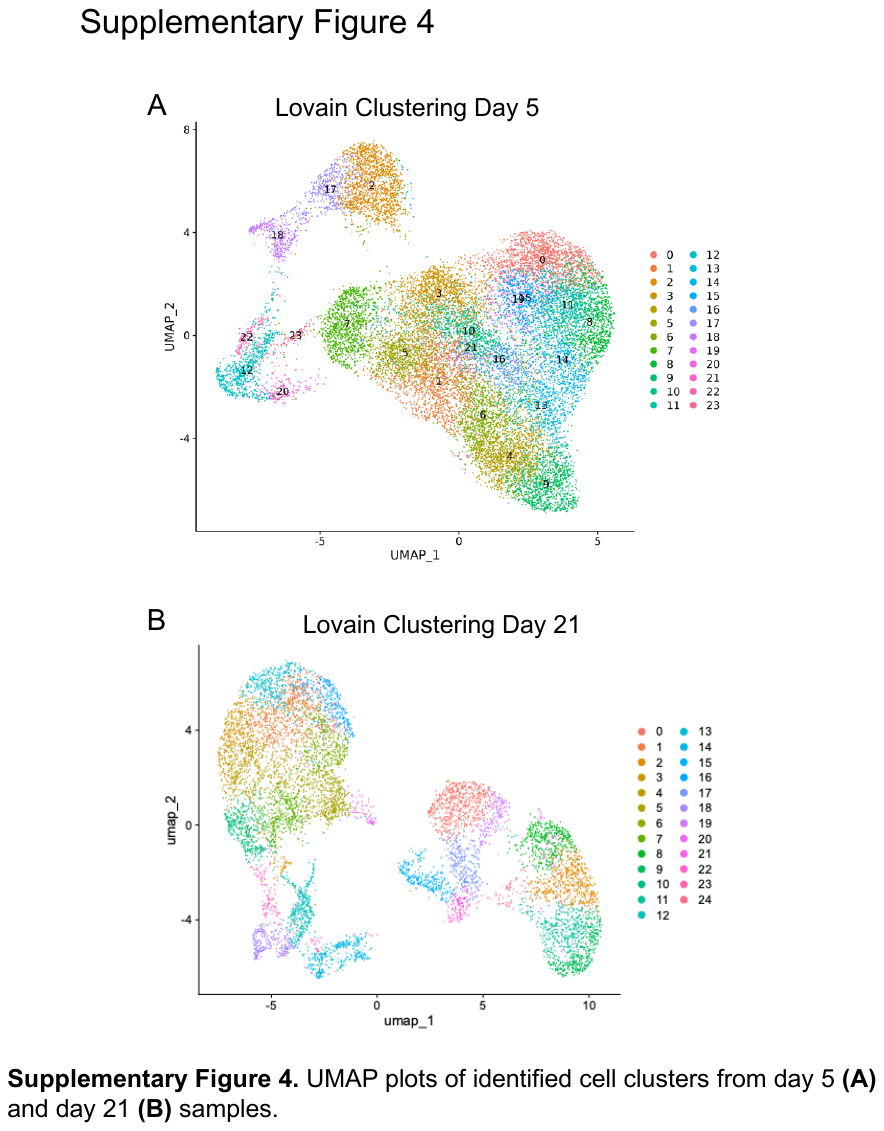

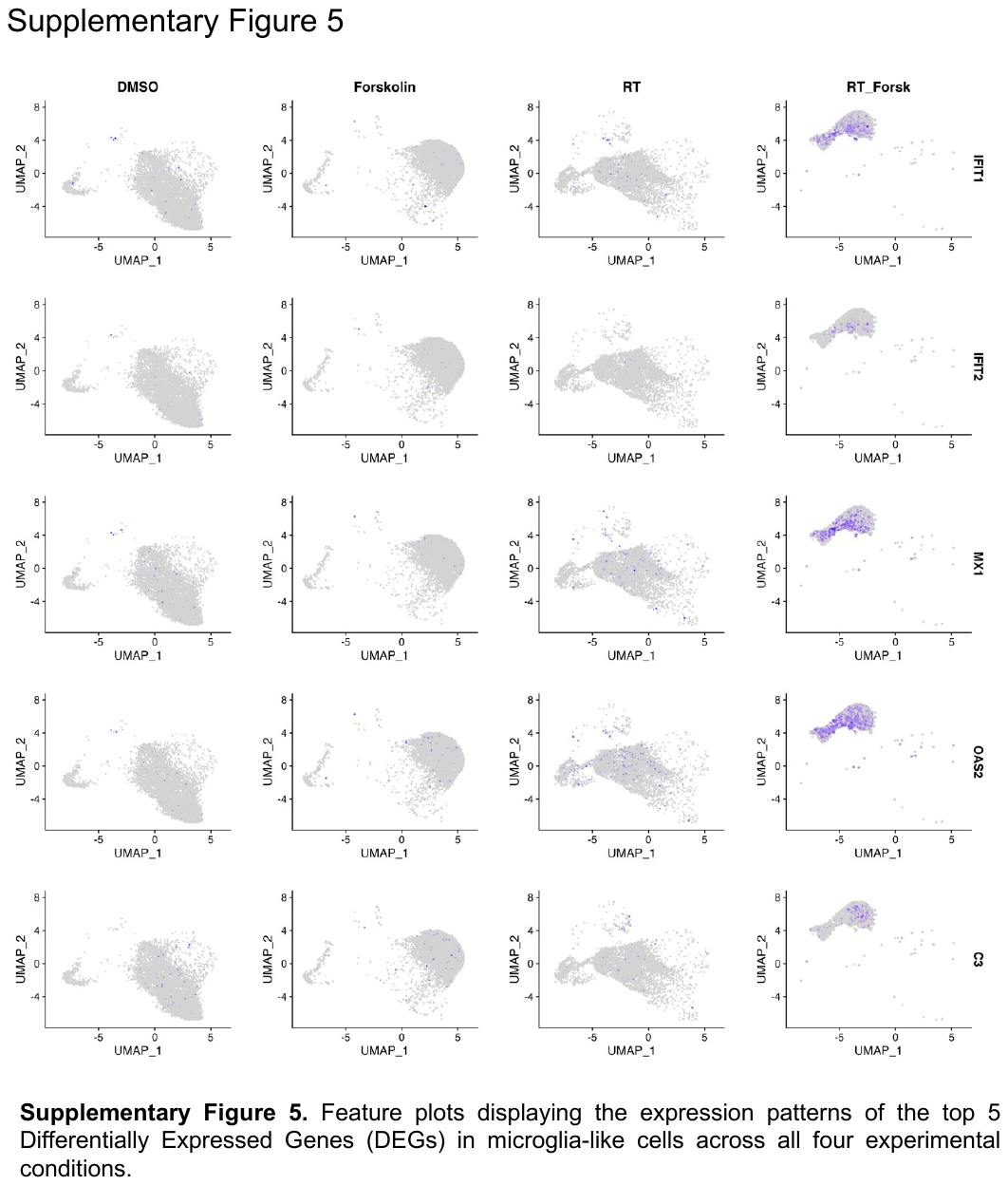

**Supplementary Tables**

**Supplementary Table 1**

| Species | Gene name | Primer sequence (5’–3’) |
| --- | --- | --- |
| Human | TUBB3 | Forward: TTTGGACATCTCTTCAGGCC  Reverse: TTTCACACTCCTTCCGCAC |
| Human | MAP2 | Forward: CTTCAGCTTGTCTCTAACCGAG  Reverse: CTGCAACTATTCAAGGAAGTGG |
| Human | MAG | Forward: CTACATTACCCAGACACGCAG  Reverse: TCTCGCTCTCGTACTTCTCTG |
| Human | CHAT | Forward: CATTGTGCAGCAGTTTGGGG  Reverse: TGCAAACCTCAGCTGGTCAT |
| Human | NF_LC | Forward: AGAGTGAAATGGCACGATACC  Reverse: ACTGGTTATGCTTCCCACG |
| Human | TMEM119 | Forward: TGTCCACCCCAGTGTCTAA  Reverse: GTGTCAGGAAGCAGTCAGG |
| Human | PPIA | Forward: ATGCTGGACCCAACACAAAT  Reverse: TCTTTCACTTTGCCAAACACC |

**Supplementary Table 2. Patient demographics and TCGA-classification of GBM subtypes.**

| **Line** | **Origin** | **Age** | **Sex** | **TCGA subtype** | **Culture P53 CN** | **EGFRvIII** | **PTEN** | **MGMT** |
| --- | --- | --- | --- | --- | --- | --- | --- | --- |
| HK-374 | Primary GBM | 45 | M | classical | Loss "mosaic" | Positive | Positive | not methylated |
| HK-157 | Primary GBM | 54 | F | proneural | wt | Negative | Positive | Unknown |
| HK-308 | Recurrent GBM | 50 | F | mesenchymal | Unknown | Positive | Positive | not methylated |

**Supplementary Table 3. Cell counts and viability of the GBM cells cultured in the Poly-D-Lysine/Laminin coated plates**

| **Experimental Group** | **Day 5** | | **Day 21** | | |
| --- | --- | --- | --- | --- | --- |
|  | **Cell counts** | **Cell viability** | **Cell counts** | | **Cell viability** |
| DMSO_1 | 2.21 x 10^6/ml | 95 % | 2.47 x 10^6/ml | 79 % | |
| DMSO_2 | 3.03 x 10^6/ml | 91 % | 1.17 x10^6/ml | 70 % | |
| DMSO_3 | 2.24 x 10^6/ml | 95 % | 2.01 x10^6/ml | 74 % | |
| Forskolin_1 | 1.48 x 10^6/ml | 89 % | 1.72 x10^6/ml | 67 % | |
| Forskolin_2 | 1.66 x 10^6/ml | 81 % | 1.41 x10^6/ml | 64 % | |
| Forskolin_3 | 1.8 x 10^6/ml | 91 % | 1.40 x 10^6/ml | 54 % | |
| RT + DMSO_1 | 9.31 x 10^5/ml | 83 % | 1.77 x 10^6/ml | 60 % | |
| RT + DMSO_2 | 1.21 x 10^6/ml | 84 % | 1.72 x10^6/ml | 62 % | |
| RT + DMSO_3 | 1.22 x 10^6/ml | 89 % | 1.77 x 10^6/ml | 67 % | |
| RT + Forskolin_1 | 3.14 x 10^5/ml | 70 % | 9.40 x 10^5/ml | 51 % | |
| RT + Forskolin_2 | 2.88 x 10^5/ml | 58 % | 1.22 x 10^6/ml | 56 % | |
| RT + Forskolin_3 | 3.82 x 10^5/ml | 68 % | 1.11 x 10^6/ml | 59 % | |

**Supplementary Table 4. Percentage Composition (Day 5) for single cell sequencing**

| **Condition** | **DMSO** | **Forskolin** | **RT** | **RT + Forskolin** |
| --- | --- | --- | --- | --- |
| Astrocyte | 0.887393 | 18.07093 | 0.705972 | 0.301594 |
| Dividing | 3.151775 | 4.522424 | 5.8195 | 16.15683 |
| G2.M | 1.499388 | 0.206418 | 6.601794 | 0.08617 |
| Glycolytic | 33.35373 | 1.332333 | 1.335623 | 0.129255 |
| Inhibitory Neuron | 15.34578 | 1.632576 | 22.76283 | 0.215424 |
| Low Quality | 0.0459 | 0.056296 | 20.56859 | 0.430849 |
| Mesenchymal | 4.023868 | 0.450366 | 5.456974 | 0 |
| Microglia | 0.0612 | 0.03753 | 0.381607 | 59.15554 |
| NPC | 15.28458 | 0.78814 | 30.24232 | 0.17234 |
| Neuron | 11.39841 | 28.33552 | 2.022515 | 22.66265 |
| OPC | 12.959 | 2.139238 | 3.415379 | 0.215424 |
| RG | 0.413097 | 4.691312 | 0.114482 | 0 |
| Vascular | 1.575887 | 37.73691 | 0.57241 | 0.473934 |

**Supplementary Table 5. Percentage Composition (Day 21) for single cell sequencing**

| **Condition** | **DMSO** | **Forskolin** | **RT** | **RT + Forskolin** |
| --- | --- | --- | --- | --- |
| Astrocyte | 12.69 | 38.22 | 12.62 | 7.59 |
| Dividing | 19.27 | 5.94 | 15.98 | 8.56 |
| G2.M | 3.9 | 0.96 | 3.05 | 1.09 |
| Glycolytic | 1.11 | 0.03 | 0.31 | 0.04 |
| Inhibitory Neuron | 3.79 | 0.2 | 3 | 0.19 |
| Low Quality | 7.57 | 1.73 | 7.12 | 14.71 |
| Mesenchymal | 25.84 | 11.38 | 26.31 | 2.41 |
| Microglia | 0.89 | 4.51 | 1.07 | 38.95 |
| NPC | 13.36 | 10.68 | 19.75 | 15.45 |
| Neuron | 10.58 | 25.28 | 9.87 | 10.23 |
| Vascular | 1 | 1.06 | 0.92 | 0.78 |

**Supplementary Table 6. Confidence intervals for stem cell frequency (%)**

| Groups | Lower | Estimate | Upper |
| --- | --- | --- | --- |
| Corn oil | 1.20772947 | 1.63934426 | 2.22222222 |
| Forskolin | 0.12487512 | 0.16207455 | 0.21052632 |
| RT + Corn oil | 0.15146925 | 0.19120459 | 0.24154589 |
| RT + Forskolin | 0.03066262 | 0.04061738 | 0.05379236 |

**Supplementary Table 7. Pairwise tests for differences in stem cell frequencies**

| Group 1 | Group 2 | Chisq | DF | Pr(>Chisq) |
| --- | --- | --- | --- | --- |
| RT + Forskolin | RT + Corn oil | 96.6 | 1 | 8.37e-23 |
| RT + Forskolin | Forskolin | 73.3 | 1 | 1.1e-17 |
| RT + Forskolin | Corn oil | 313 | 1 | 5.7e-70 |
| RT + Corn oil | Forskolin | 1.44 | 1 | 0.23 |
| RT + Corn oil | Corn oil | 125 | 1 | 5.19e-29 |
| Forskolin | Corn oil | 142 | 1 | 7.84e-33 |
